# NGFR-driven suppression of antigen presentation limits CD8^+^ T cell immunity and response to checkpoint blockade

**DOI:** 10.64898/2026.08.24.746602

**Authors:** Juan García-Agulló, Vanesa Santos, Blanca Majem, Mariola Munárriz, Laura Serrano, María Calvo de Mora, Sara Sánchez-Redondo, Daniel Achuela, Teresa Aceña-Gonzalo, Inés Sentís, Gloria Pascual, Carmen Blanco-Aparicio, Fátima Al-Shahrour, Eduardo Caleiras, Isabel Peset, Juan P. Rodrigo, Juana M. García-Pedrero, Mónica Álvarez-Fernández, H. Uri Saragovi, Laura Nogués, María Casanova-Acebes, Salvador Aznar-Benitah, Héctor Peinado

## Abstract

Immune checkpoint blockade has revolutionized cancer therapy; however, numerous tumors remain resistant by adopting cellular states that impede immune recognition. In this study, we identify the nerve growth factor receptor (NGFR) as a regulator of immune evasion in head and neck squamous cell carcinoma (HNSCC). Genetic ablation of *Ngfr* resulted in impaired tumor growth in immunocompetent MOC2 HNSCC, while pharmacological inhibition with THX-B reduced primary tumor growth and spontaneous metastatic dissemination. Single-cell profiling of MOC2 tumors demonstrated that *Ngfr* loss redirected tumor cells away from invasive EMT-like states and enhanced antigen-processing and presentation programs. This was accompanied by increased presentation of tumor antigens and expansion of effector CD8⁺ T-cells *in vivo*. Functionally, CD8⁺ T-cell depletion, Batf3 deficiency, and JAK1/2 inhibition restored the growth of *Ngfr*-deficient tumors, indicating that NGFR loss exposes tumors to CD8⁺ T-cell–mediated control through a JAK-associated antigen-presentation program. Notably, NGFR blockade sensitized otherwise resistant MOC2 tumors to anti-PD1 therapy, and the combination of THX-B with anti-PD1 significantly improved tumor control and survival. In human HNSCC, spatial profiling revealed that NGFR⁺ tumor regions exhibited reduced HLA-DR expression and limited CD3⁺ T-cell infiltration. Notably, an NGFR-associated antigen-presentation signature stratified survival and response in HNSCC patients undergoing immune checkpoint blockade. Interestingly, this signature was also linked to improved outcomes in melanoma patients. We also observed a significant increase in the effector CD8⁺ T-cell fraction in melanoma NGFR KO tumors linked to a significant decrease in tumor growth. These findings position NGFR as a regulator of tumor immune visibility and support NGFR inhibition as a strategy to enhance immunotherapy response.

## Introduction

The nerve growth factor receptor (NGFR), also known as CD271 or p75^NTR^, is a low-affinity transmembrane receptor for neurotrophins and proneurotrophins that plays a central role in regulating cellular plasticity in cancer [1]. Traditionally, NGFR has been described as a modulator of central nervous system function, controlling processes such as survival, apoptosis, cell proliferation, and migration [1]. In cancer, NGFR was first identified as a marker of tumor cells with stem-like traits across different tumors, including melanoma [2–4], breast cancer [5], colon cancer [6], esophageal cancer [7], and HNSCC [8, 9]. In HNSCC, in particular, NGFR has been associated with poor prognosis [10], high tumorigenic capacity *in vivo* [8], metastatic potential [11], and a highly invasive, epithelial-mesenchymal transition (EMT)-like phenotype [11, 12].

Importantly, NGFR is associated with the immune-evasive properties of melanoma. NGFR expression is inversely correlated with the expression of MHC class I molecules and other immune ligands, such as NK-activating ligands CD112 and CD155, suggesting that NGFR reduces tumor immunogenicity [13–15]. In addition, NGFR expression is associated with transcriptional signatures of resistance to immunotherapy [14]. In melanoma, longitudinal analysis of delayed resistance to immune checkpoint blockade revealed the emergence of NGFR-expressing tumor cell states in relapsing lesions positioned in close proximity to immune cells. These data suggest a link between NGFR⁺ tumor states and immune escape [16]. More recently, pharmacological inhibition of NGFR using THX-B has been shown to reduce metastatic dissemination, increase intratumoral CD8⁺ T-cell infiltration, and enhance the efficacy of immune checkpoint blockade in immunotherapy-resistant melanoma models [17]. However, the molecular mechanisms by which NGFR orchestrates tumor immune evasion, and whether this immunosuppressive axis operates across tumor types, remain incompletely understood.

Immunotherapy, particularly immune checkpoint blockade (ICB), has transformed cancer treatment, especially for advanced and metastatic diseases. Several clinical trials have demonstrated its efficacy in different tumor types, including head and neck squamous cell carcinoma (HNSCC) [18–20], melanoma [21, 22], non-small cell lung cancer [23–25], and triple-negative breast cancer [26, 27], among others. However, PD-1 inhibitors, the cornerstone of ICB, provide significant clinical benefit to only a minority of patients, with objective response rates typically ranging from approximately 10–25% [28]. Therefore, understanding the immune evasion mechanisms that result in primary and acquired resistance to ICB remains a priority in cancer research.

In this study, we identify NGFR as a regulator of immune evasion in HNSCC and melanoma. Using immunocompetent models, we show that genetic deletion or pharmacological inhibition of NGFR restrains primary tumor growth and spontaneous metastatic dissemination. In HNSCC, NGFR loss reshaped malignant cell states, reducing invasive/EMT-like programs while inducing an inflammatory antigen-processing and presentation phenotype. This remodeling increased tumor antigen presentation promoted cytotoxic effector CD8⁺ T-cell expansion and enhanced the accumulation of antigen-specific T cells in vivo. The antitumor effect of NGFR loss was reversed by CD8⁺ T-cell depletion, Batf3 deficiency or JAK inhibition, indicating that NGFR limits a JAK-associated antigen-presentation program required for productive CD8⁺ T-cell immunity. Consistently, pharmacological NGFR blockade sensitized resistant tumors to anti-PD1 therapy. In human HNSCC, NGFR⁺ tumor regions displayed reduced HLA-DR expression and poor CD3⁺ T-cell infiltration, while diffuse NGFR expression was associated with aggressive and metastatic disease. Moreover, an NGFR-associated antigen-presentation signature stratified survival and response in HNSCC patients receiving immune checkpoint blockade and was linked to improved outcome in melanoma patients. Together, these findings establish NGFR as a suppressor of tumor immune visibility and support NGFR blockade, together with NGFR-derived signatures, as complementary therapeutic and biomarker strategies to enhance immunotherapy response.

## Results

### NGFR blockade impairs tumor progression in immunocompetent models of HNSCC

HNSCC is frequently presented at advanced stages with poor prognosis and notable resistance to immunotherapy [18–20]. While NGFR has been associated with aggressive phenotypes in HNSCC [12], its role in immune modulation in this tumor type remains unclear. To investigate the contribution of NGFR to tumor growth and immune resistance in HNSCC, we selected the MOC2 model [29]. This immunocompetent murine model forms tumors *in vivo* [29, 30] and reproduces features of aggressive disease, including an immunosuppressive and immunotherapy-resistant tumor microenvironment (TME) [31, 32].

To evaluate the impact of *Ngfr* loss on tumor growth *in vivo*, we generated an *Ngfr*-knockout (Ngfr KO) MOC2 model and orthotopically implanted these cells into C57BL/6 mice, monitoring tumor progression longitudinally by bioluminescence imaging (BLI)). *Ngfr* knockout efficiency was confirmed both *in vitro* by immunoblotting (**Fig. 1a**) and *in vivo* by immunohistochemistry (IHC) in tumors (**Extended Data Fig. 1a**). On day 13 post-injection mice bearing MOC2 GL *Ngfr*-KO tumors displayed a significantly lower tumor burden than mice bearing MOC2 GL control tumors (**Fig. 1b-d**, Extended Data Fig. 1b).

**Fig. 1.**
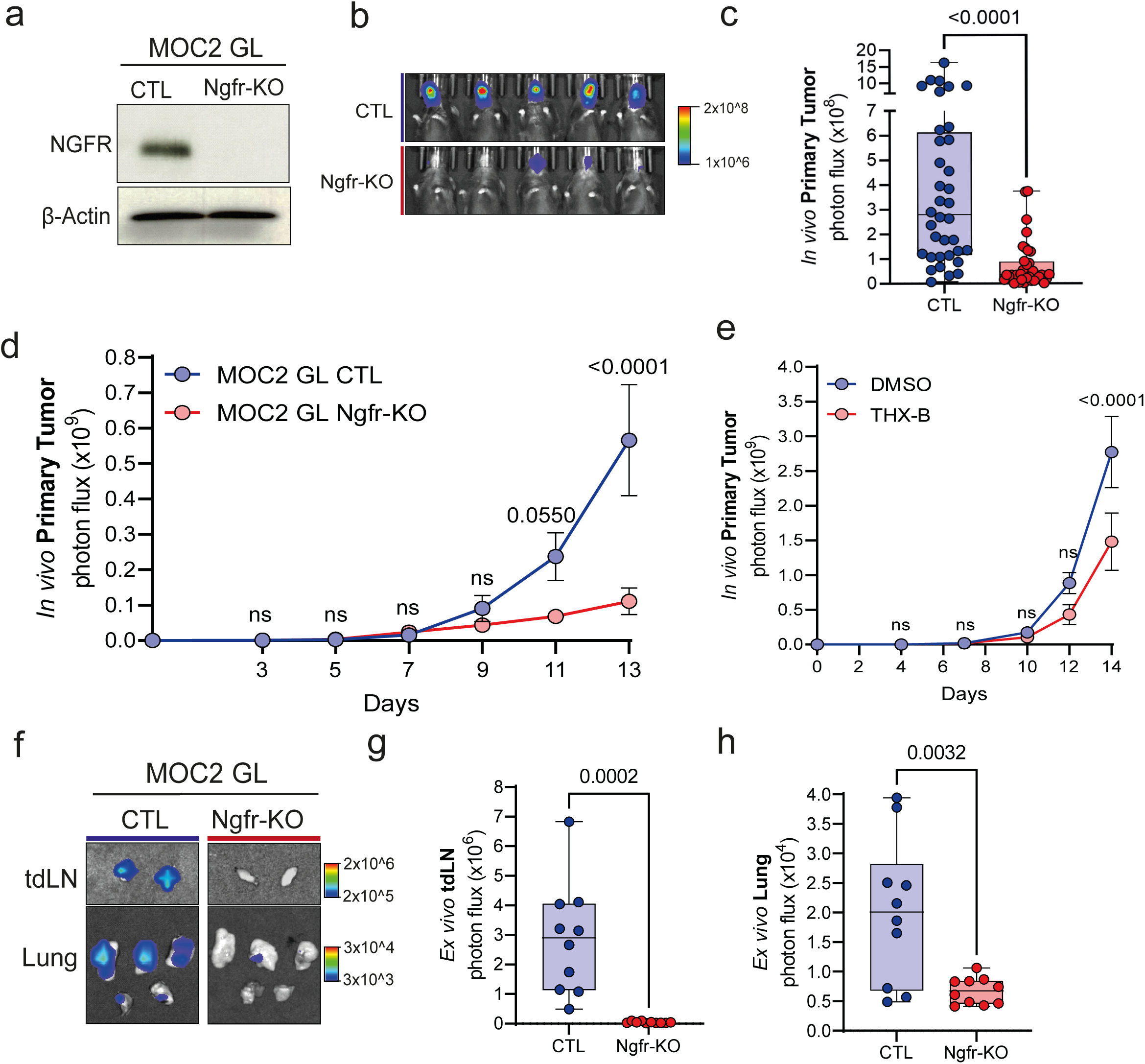
| NGFR blockade in an immunocompetent setting impairs HNSCC primary tumor growth. a, Validation of Ngfr-KO in vitro by immunoblotting (n = 3). b, c, Tumor burden at experimental endpoint (D13) comparing MOC2 GL CTL and Ngfr-KO tumors. Representative bioluminescence images from one experiment are shown in b, along with a summary box plot of all experiments in c. Each dot represents an individual mouse. Statistical analysis was performed using an unpaired t-test. Sample size: n = 36 mice per group, pooled from five independent experiments. d, Longitudinal tumor growth analysis based on in vivo BLI at D3, 5, 7, 9, 11, and 13, showing the mean tumor growth per group ± SEM for MOC2 GL CTL and Ngfr-KO tumors. Statistical comparisons at each time point were performed using two-way ANOVA (ns = not significant). Sample size: n = 11 for MOC2 GL CTL–bearing mice and n = 13 for MOC2 GL Ngfr-KO–bearing mice, pooled from two experiments. e, Longitudinal tumor growth comparison between DMSO and THX-B treatment groups, showing the mean ± SEM at each time point. Statistical comparisons at each time point were performed using two-way ANOVA. Sample size: n = 19 for DMSO-treated, and n = 18 for THX-B-treated mice, pooled from two independent experiments. f, Representative tdLN and lung metastases from an individual mouse from the CTL or Ngfr-KO groups. g, h, Quantification of metastases. tdLN metastasis is shown in g, and lung metastasis is shown in h. Each dot represents one mouse; tdLN and lung signals represent the combined signal from both cervical lymph nodes and all five lung lobes, respectively. Sample size: n = 10 mice per group, pooled from two independent experiments.

We next explored a pharmacological approach to inhibit NGFR using THX-B, a small molecule inhibitor reported to block NGFR [33] and reduce metastasis in melanoma models [17, 34]. MOC2 GL cells were orthotopically implanted, and mice were treated with THX-B or vehicle starting on day 6 (D6), with tumor growth monitored until day 14. THX-B treatment significantly reduced primary tumor growth (**Fig. 1e**, Extended Data Fig. 1c), further supporting the therapeutic potential of NGFR inhibition in oral cancer.

In addition to its effects on primary tumor growth, we evaluated whether NGFR also contributes to spontaneous metastatic dissemination in the orthotopic MOC2 GL model. We observed that *Ngfr*-KO significantly reduced metastasis to tumor-draining cervical lymph nodes (tdLNs) (**Fig. 1f, g)** and lungs **(Fig. 1f, h).** Similarly, pharmacological inhibition with THX-B decreased metastatic dissemination to tdLNs **(Extended Data Fig. 1d)** and lungs **(Extended Data Fig. 1e**). These findings are consistent with previous melanoma studies showing that THX-B limits NGFR-dependent metastatic dissemination to lymph nodes and lungs [34].

To investigate whether the effect of *Ngfr* loss on metastatic dissemination depended on the orthotopic primary tumor setting, we performed an experimental metastasis assay by tail vein injection of MOC2 GL control or *Ngfr*-KO cells into C57BL/6 mice. In contrast to the orthotopic model, ex vivo BLI detected no significant differences in lung metastatic burden between the groups **(Extended Data Fig. 1f, g**).

### NGFR depletion reprograms tumor cell states and enhances antigen presentation

To assess the molecular changes induced by *Ngfr* loss, we performed single-cell RNA sequencing (scRNA-seq) of equal numbers of tumor (GFP⁺) cells isolated from orthotopic MOC2 GL CTL and Ngfr-KO tumors. To account for the tumor growth differences already apparent at day 13, equal numbers of both malignant (GFP⁺) and immune (CD45⁺) cells were processed per condition across all scRNA-seq experiments, ensuring that transcriptional profiles reflect Ngfr-dependent reprogramming rather than differences in tumor cellularity. Unsupervised analysis of tumor cells revealed multiple genes involved in the regulation of the epithelial cell phenotype or epithelial-to-mesenchymal transition (EMT) (**Extended Data Fig. 2a, b**). We found that *Ngfr*-KO significantly reduced late EMT clusters: the “EMT late/invasive” cluster by more than 4-fold, and the “EMT late/contractile” cluster was reduced by more than 2.5-fold (**Fig. 2a, c and Extended Data Fig. 2c**). In contrast, epithelial clusters were increased in Ngfr-KO tumors, with a 1.4-fold increase in the “Epithelial differentiated” cluster and a 1.8-fold increase in the “Epithelial proliferative” cluster **(Fig. 2b, c** and **Extended Data Fig. 2c**). These results indicate that NGFR in tumor cells favors an EMT-like transcriptional program in HNSCC, in line with previous studies in different tumors [2–9]. Moreover, these results suggest that the NGFR-driven EMT-like phenotype is reversible and shifts toward a more epithelial state upon *Ngfr* depletion.

**Fig. 2.**
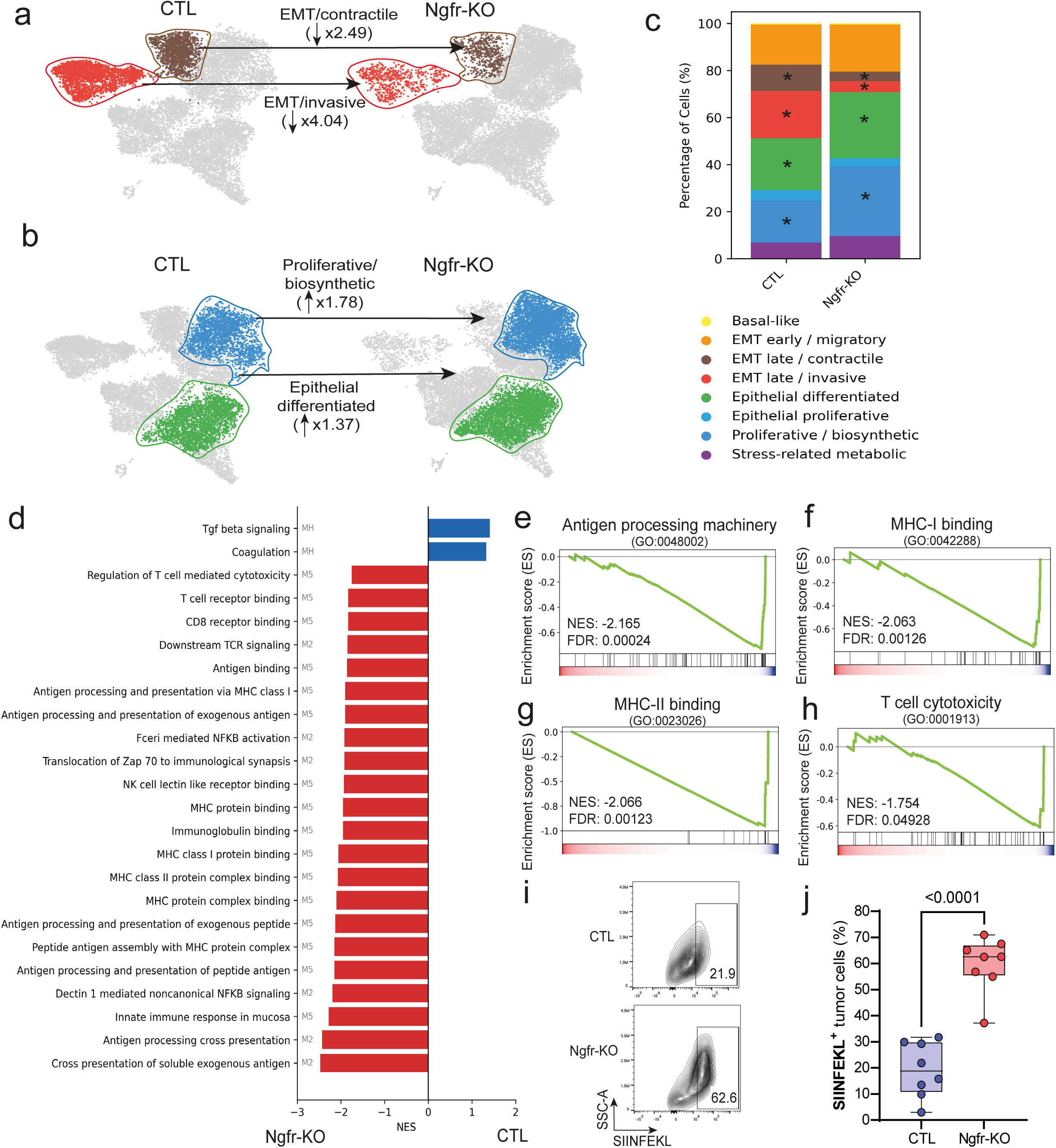
| NGFR suppresses EMT and antigen presentation programs in cancer cells. a, b, UMAP representations of tumor cells from MOC2 GL CTL (left) and Ngfr-KO (right) tumors, highlighting the EMT late clusters (“EMT late/invasive” in red and “EMT late/contractile” in brown) in a, and the “Epithelial differentiated” (green) and “Proliferative/biosynthetic” (blue) clusters in b. Numbers in parentheses indicate the fold change in cluster abundance in Ngfr-KO relative to CTL. c, Stacked bar plots showing the relative percentage of each tumor cluster in MOC2 GL CTL or Ngfr-KO samples. Statistical significance of changes in cluster proportions (denoted by an asterisk) was determined using a two-step approach: (I) a contingency table test (Fisher’s exact test for cell counts <5, or chi-square test otherwise) with FDR adjustment (FDR < 0.05), and (II) biological relevance, defined as an absolute percentage change greater than 5%. d, GSEA of all significantly dysregulated immune pathways (FDR < 0.05) comparing MOC2 CTL and Ngfr-KO tumor cells. Hallmarks (MH), Reactome (M2), and Gene Ontology (M5) mouse datasets were used and combined in a single graph, with the source dataset indicated to the left of each pathway name. FDR values were calculated using the Benjamini-Hochberg method to adjust for multiple comparisons. e-h, GSEA enrichment plots of key immune signatures upregulated in MOC2 GL Ngfr-KO tumor cells, including antigen processing and presentation in e, MHC class I in f, MHC class II protein binding in g, and T cell cytotoxicity regulation in h. FDR, NES and the corresponding GO term for each GSEA enrichment plot are shown. i, Representative density plots of SIINFEKL membrane expression in one mouse from each group. j, Quantification of the percentage of SIINFEKL+ tumor cells in MOC2 GL CTL or Ngfr-KO tumors. Statistical comparison was performed using an unpaired t-test. Sample size: n = 8 mice per group, from two independent experiments.

To identify NGFR-regulated pathways, we performed differential expression analysis between MOC2 GL CTL and *Ngfr*-KO tumor cells, followed by gene set enrichment analysis (GSEA). Interestingly, we observed significant enrichment of multiple antigen-processing machinery (APM)-related pathways in Ngfr-KO cells (**Fig. 2d**). Among these pathways we found a significant enrichment of antigen processing machinery (GO: 0048002) (**Fig. 2e**), MHC class I-mediated antigenic presentation (GO: 0042288) (**Fig. 2f**), binding to MHC class II complex (GO: 0023026) (**Fig. 2g**), and T-lymphocyte-mediated cytotoxicity (GO: 0001913) (**Fig. 2h**), suggesting that NGFR is controlling antigen presentation and T-Cell-mediated surveillance in HNSCC.

To analyze the impact of NGFR on antigen presentation, we generated MOC2 GL CTL and Ngfr-KO lines expressing OVA-mCherry and orthotopically injected them into C57BL/6 mice. In this model, *Ngfr*-KO tumor cells showed increased SIINFEKL peptide expression at the plasma membrane (**Fig. 2i-j**). To analyze whether this effect was due to differences in tumor size at the endpoint, we analyzed SIINFEKL presentation at an earlier time point (day 9; D9) when no significant differences in tumor size were detected (**Extended Data Fig. 2d**). Remarkably, we observed that Ngfr KO promoted the presentation of SIINFEKL peptide from this early timepoint (**Extended Data Fig. 2e**).

### NGFR blockade expands CD8^+^ T effector cells impairing primary tumor growth

Given the increased antigen presentation observed in Ngfr-KO tumor cells, we examined whether NGFR influences the immune landscape of the TME. We performed scRNA-seq analysis of CD45^+^ immune cells isolated from MOC2 GL CTL and *Ngfr*-KO. Major immune cell populations were identified and annotated based on canonical markers (**Extended Data Fig. 3a, b**). The myeloid compartment was dominated by neutrophils (S100a9^+^) and macrophages (Fcgr1^+^), with smaller populations of monocytes (Fcgr3^+^), dendritic cells (Flt3^+^), and mast cells (Fcerc1a^+^) present at lower frequencies (**Extended Data Fig. 3b-d**). Lymphoid compartment was enriched in T cells (Cd3d/e^+^) followed by NK cells (Klrb1c^+^) (**Extended Data Fig. 3b, e, f**). The absolute numbers and relative abundance of each immune population in MOC2 GL CTL and Ngfr-KO tumors are shown in **Extended Data Fig. 3g**.

We next examined the T cell compartment in greater detail. Flow cytometry analyses confirmed no significant differences in the frequency of total T cells, CD4⁺ T cells, or CD8⁺ T cells (Extended Data Fig. 4a**–e**). We therefore asked whether NGFR affected the distribution of specific T cell states. Unsupervised subclustering of the scRNA-seq dataset identified ten T cell clusters (**Fig. 3a-c, Extended Data Fig. 4f**), including four CD4^+^ subtypes, three CD8^+^ clusters, two mixed CD4^+^/CD8^+^ populations, and one CD4^-^/CD8^-^ Th17 population. Among these, MOC2 GL Ngfr-KO tumors showed marked (>2.5-fold) expansion of CD8^+^ effector cytotoxic T cells (**Fig. 3d, e**) defined by the expression of classical cytotoxic markers, such as *Prf1* or *Gzmb* (**Extended Data Fig. 4g, h**). Importantly, Ngfr-KO tumors also exhibited a reduction in the CD8+ exhausted T cell cluster (**Extended Data Fig. 4f**), indicating that effector cells remain functionally active and are not terminally exhausted, a phenotypic distinction relevant to therapy-resistance contexts. This increase in effector cells was validated by flow cytometry as an expansion of CD8^+^ effector T cells (CD44^+^, CD62L^-^) in MOC2 GL Ngfr-KO tumors (**Fig. 3f, g**) and was also observed after THX-B treatment (**Extended Data Fig. 4i**).

**Fig. 3.**
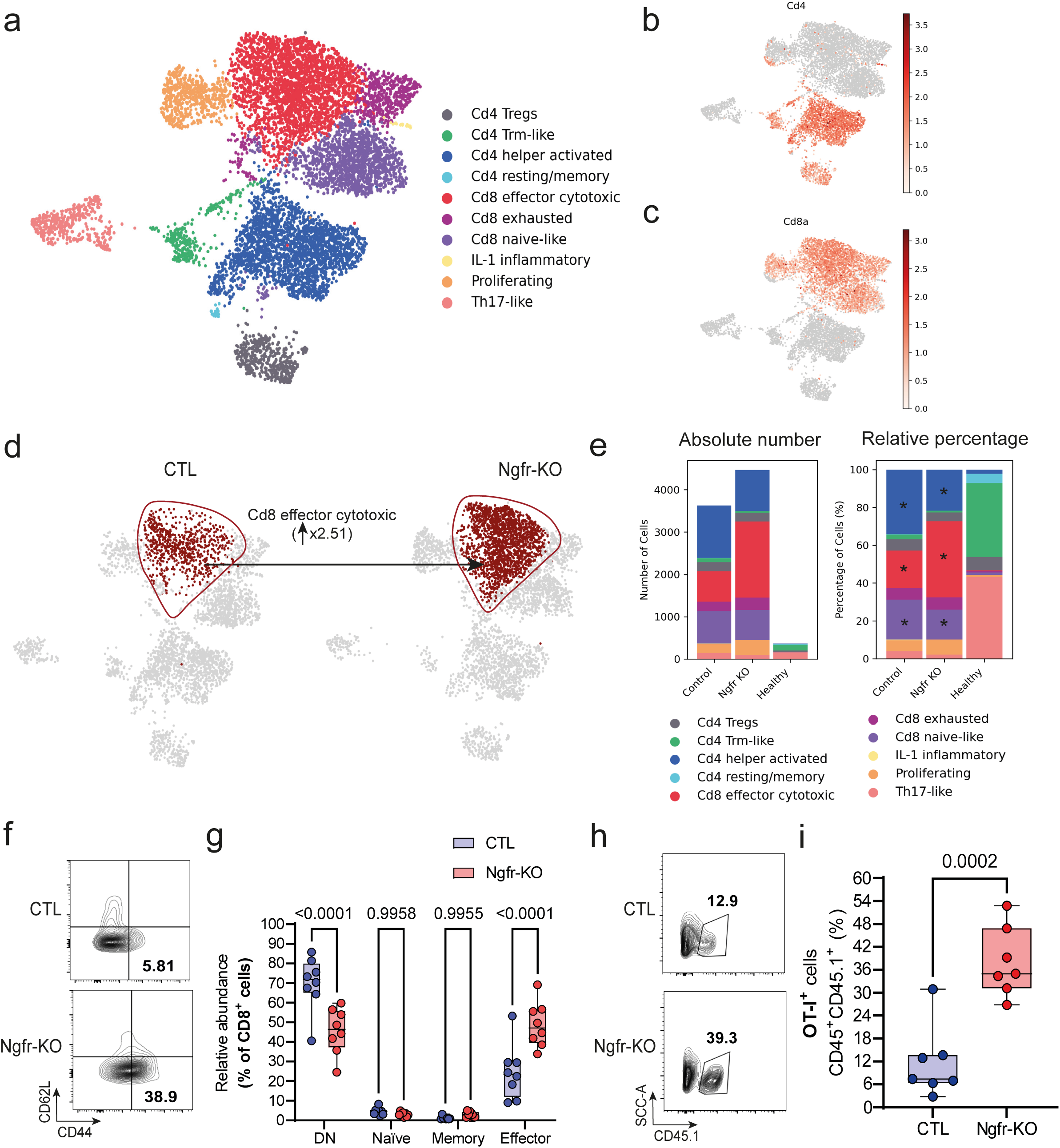
| NGFR depletion expands CD8+ effector T cells. a, UMAP projection of T cell subcluster annotations based on unbiased clustering of CD3+ T cells performed with Scanpy. b, c, UMAP representations of Cd4 (b) and Cd8a (c) expression across T cell subclusters. d, UMAP highlighting the CD8⁺ effector cytotoxic T cell population in MOC2 GL CTL and Ngfr-KO tumors (fold change = 2.51; highlighted in red). e, Stacked bar plots showing the absolute number of cells (left) and the relative percentage (right) of each T cell subcluster in MOC2 GL CTL and Ngfr-KO tumors, together with healthy (non-tumor-bearing) tissue as a reference. Statistical significance (denoted by an asterisk) was determined using a two-step approach: (I) a contingency table test (Fisher’s exact test for cell counts <5 or chi-square test otherwise) with FDR adjustment (FDR < 0.05), and (II) biological relevance defined as an absolute percentage change greater than 5%. f, Density plots representative of CD8+ T cell (CD45+, CD3+, CD8+) subtypes in MOC2 GL CTL or Ngfr-KO tumors. Subtypes were defined as: naïve (CD44−, CD62L+), central memory (CD44+, CD62L+), effector (CD44+, CD62L−), and double negative (DN; CD44−, CD62L−). g, Quantification of the percentages of each subtype relative to total CD8+ T cells in MOC2 GL CTL or Ngfr-KO tumors. P values were calculated using two-way ANOVA. Sample size: n = 8 mice per group, combined from two independent experiments. h, Density plots representative of OT-I cells (CD45+, CD45.1+) in tumors derived from MOC2 GL OVA-mCherry CTL or Ngfr-KO at D14 after tumor injection. i, Quantification of the percentages of OT-I cells relative to total CD45+ cells in MOC2 GL OVA-mCherry CTL or Ngfr-KO tumors. P value was calculated using an unpaired t-test. Sample size: n = 7 mice per group, combined from two independent experiments.

To examine whether the increased CD8+ effector T cell response in Ngfr-KO tumors was associated with tumor antigen presentation, we used the OVA-mCherry model. MOC2 CTL OVA-mCherry CTL or *Ngfr*-KO tumor cells were implanted into C57BL/6 mice, followed by adoptive transfer of OT-I/CD45.1 T cells after tumor establishment. Consistent with previous results, Ngfr-KO primary tumors showed reduced growth compared to CTL (**Extended Data Fig. 4j, k)**. At the endpoint, Ngfr-KO tumors contained higher numbers of OVA-specific OT-I cells (CD45^+^CD45.1^+^) (**Fig. 3h, i**), which were validated as CD3e^+^CD8a^+^SIINFEKL-tetramer^+^ cells (**Extended Data Fig. 4l**).

The expansion of effector CD8^+^ T cells observed in MOC2 *Ngfr*-KO tumors prompted us to test whether these cells contribute to the reduced tumor growth phenotype. CD8^+^ T cells were depleted *in vivo* using an anti-CD8 monoclonal antibody. We confirmed the depletion of CD8+ T cells after treatment by flow cytometry (**Fig. 4a).** In this setting, the growth defect of MOC2 GL Ngfr-KO tumors was restored **(Fig. 4b, c**). We next tested this in Batf3^-/-^ mice^88^, which lack conventional type 1 dendritic cells and have impaired tumor antigen cross-presentation, resulting in reduced cytotoxic CD8+ T cell responses. Consistent with this model, CD8+ T cell abundance was reduced (**Fig. 4d)**, and under these conditions, MOC2 Ngfr-KO tumors showed restored growth comparable to that of the controls **(Fig. 4e, f**).

**Fig. 4.**
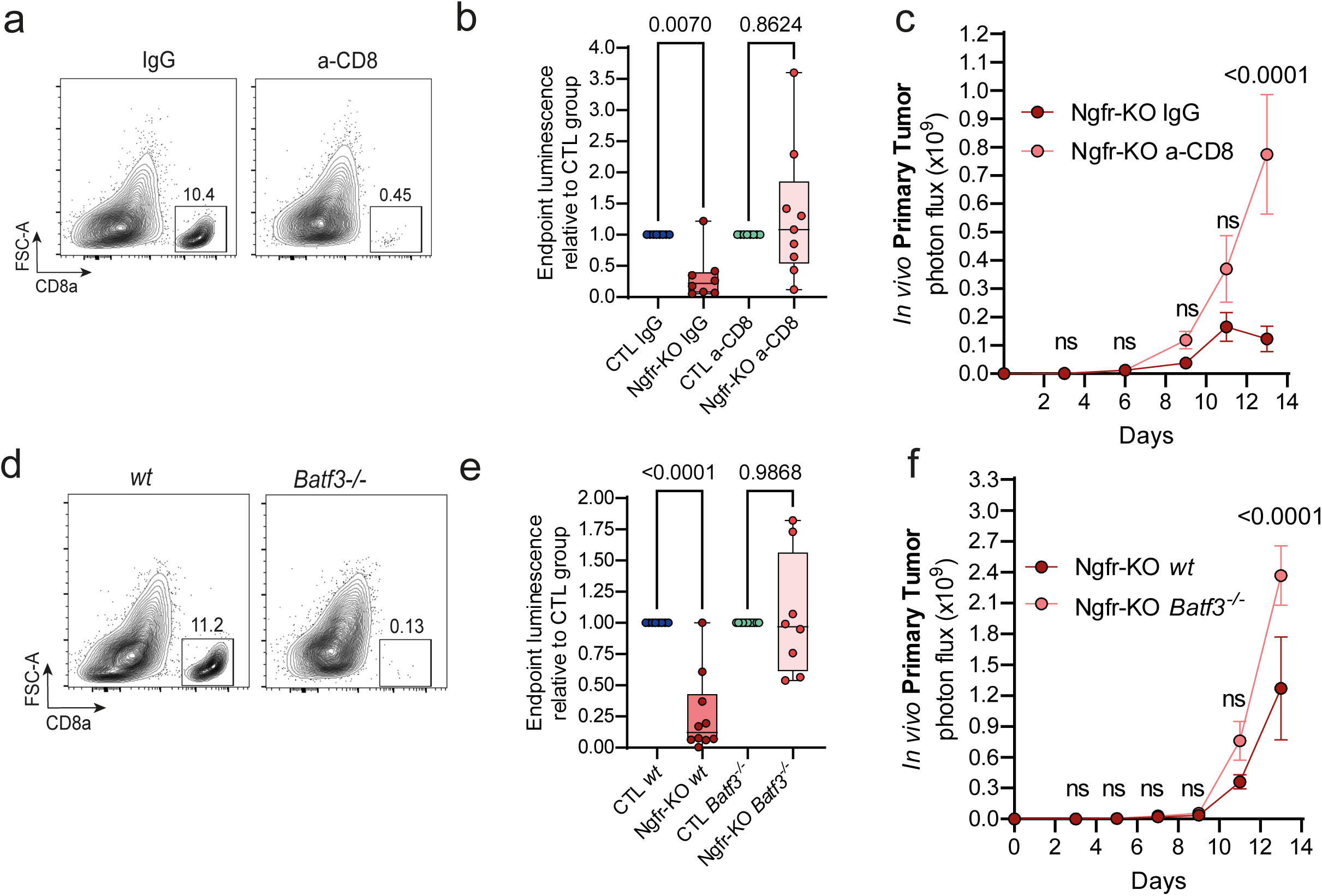
| Depletion of CD8⁺ T cells reverses tumor growth control in Ngfr-KO tumors. a, Flow cytometry density plots of CD8⁺ T cells in IgG- or anti-CD8-treated MOC2 GL CTL or Ngfr-KO tumors; the percentage of CD8⁺ cells relative to total CD45⁺ cells is shown. b, Box plot showing the primary tumor BLI signal on D13 post-injection in mice bearing MOC2 GL CTL or Ngfr-KO tumors treated with IgG or anti-CD8 antibody, expressed as a relative change compared to the control group. P values were calculated using an unpaired t-test. c, Tumor growth curves comparing MOC2 GL Ngfr-KO tumors in mice treated with IgG or anti-CD8, grouped by condition, showing the mean ± SEM. Sample size: 8 mice in each group of CTL IgG, Ngfr-KO IgG, and CTL anti-CD8; 9 mice in the Ngfr-KO anti-CD8 group, combined from two independent experiments. Statistical comparisons were performed using two-way ANOVA, comparing each time point separately. d, Flow cytometry density plots confirming the loss of the CD8α⁺ dendritic cell compartment in Batf3⁻/⁻ mice compared with wt mice; the percentage of CD8α⁺ cells out of total CD45⁺ cells is shown. e, Box plot showing the primary tumor luminescence signal on D13 post-injection in Ngfr-KO tumors injected into WT or Batf3⁻/⁻ mice, expressed as a relative change compared to their corresponding control group. P values were calculated by unpaired t-test. f, Tumor growth curves comparing MOC2 GL Ngfr-KO tumors in WT or Batf3⁻/⁻ mice, grouped by condition, showing mean ± SEM. Sample size: 8 mice in WT CTL, 10 in WT Ngfr-KO, 11 in Batf3⁻/⁻ CTL, and 8 in Batf3⁻/⁻ Ngfr-KO, combined from two independent experiments. Statistical comparisons were calculated by two-way ANOVA, comparing each time point separately.

### NGFR restrains a JAK-dependent antigen-presentation program that supports CD8⁺ T-cell immunity *in vivo*

To investigate how NGFR loss promotes antigen presentation and CD8⁺ T-cell immunity *in vivo*, we interrogated immune-related pathways in malignant cells from the MOC2 GL CTL and Ngfr-KO scRNA-seq dataset. Among the pathways enriched upon Ngfr depletion, the IFN-γ response emerged together with antigen-processing and presentation programs, including genes involved in JAK/STAT signaling, immunoproteasome activity and peptide transport, such as Jak2, Stat1, Irf1, Psmb family members and Tap transporters (**Fig. 5a**).

**Fig. 5.**
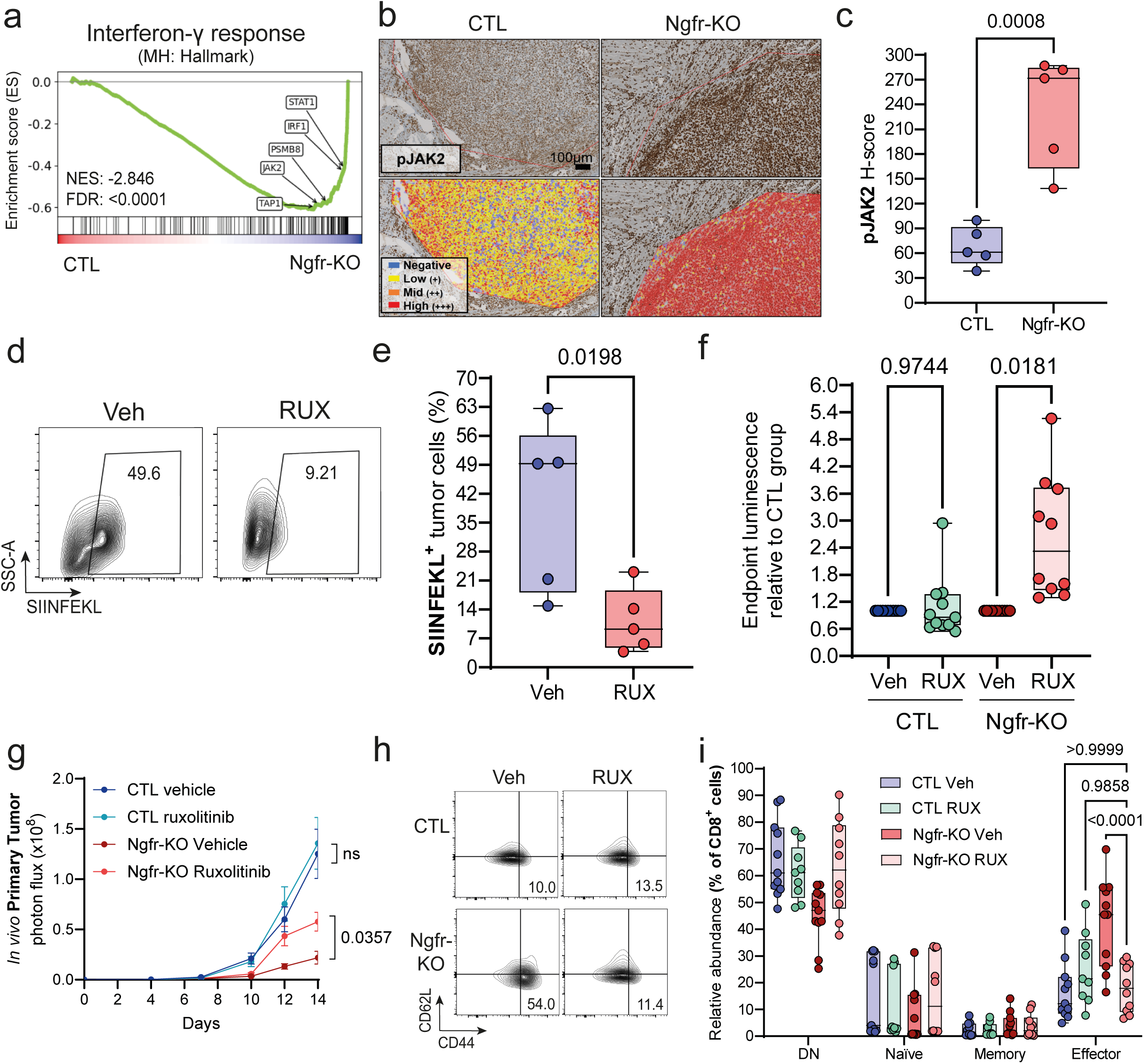
| NGFR regulates antigen presentation through JAK signaling. a, GSEA plot derived from the scRNA-seq dataset of MOC2 GL tumor cells (GFP^+^). Comparison between MOC2 GL CTL and Ngfr-KO cells, highlighting enrichment of an IFN-γ signature. JAK-related antigen presentation genes are indicated (JAK2, STAT1, IRF1, PSMB8 and TAP1). FDR values were calculated using the Benjamini-Hochberg method to adjust for multiple comparisons. b, c, Immunohistochemistry analysis of pJAK2 expression in MOC2 GL CTL or Ngfr-KO orthotopic tumors. Representative images are shown in panel b, with pJAK2 original staining (top) and quantification using QuPath based on staining intensity levels (bottom). Blue is negative, yellow is low (+), orange is intermediate (++) and red is high (+++). Quantifications of tumor H-score (calculated as intensity low × 1 + intermediate × 2 + high × 3) are shown in panel c. P value was calculated by unpaired t-test. Sample size: n = 5 per group from one experiment. d, Representative density plots of SIINFEKL membrane expression in MOC2 GL OVA-mCherry tumor-bearing mice treated with vehicle (CMC-Na) or RUX. e, Quantification of the percentage of SIINFEKL+ tumor cells in MOC2 GL OVA-mCherry vehicle- or RUX - treated mice. Statistical comparison was performed using an unpaired t-test. Sample size: n = 5 mice per group, from one experiment. f, Box plot showing the primary tumor BLI signal on D14 post-injection in mice bearing MOC2 GL CTL or Ngfr-KO tumors treated with vehicle or RUX, expressed as a relative change compared to the control group. P values were calculated by unpaired t-test. g, Longitudinal tumor growth curves determined by BLI in mice bearing MOC2 GL CTL or Ngfr-KO tumors treated with vehicle or RUX, presented as mean ± SEM. Statistical comparisons were performed at each time point using two-way ANOVA (ns = not significant). Sample sizes: n = 11 mice per group, from two independent experiments. h, Density plots representative of CD8+ T cell (CD45+, CD3+, CD8+) subtypes in MOC2 GL CTL or Ngfr-KO tumors treated with vehicle or RUX. Subtypes were defined as: naïve (CD44−, CD62L+), central memory (CD44+, CD62L+), effector (CD44+, CD62L−), and double negative (DN; CD44−, CD62L−). The percentages indicated correspond to the effector CD8+ T cells of the representative sample. i, Quantification of the percentages of each subtype relative to total CD8+ T cells in MOC2 GL CTL or Ngfr-KO tumors treated with vehicle or RUX. P values were calculated using two-way ANOVA. Sample sizes: CTL vehicle (n = 11), CTL RUX (n = 9), Ngfr-KO vehicle (n = 11), Ngfr-KO RUX (n = 10), from two independent experiments.

Consistent with this transcriptional state, MOC2 GL Ngfr-KO tumors showed increased pJAK2 levels compared with CTL tumors, supporting enhanced activation of a JAK-associated program *in vivo* (**Fig. 5b-c**). To determine whether JAK signaling was required for the enhanced antigen presentation induced by NGFR loss, we treated MOC2 GL OVA-mCherry Ngfr-KO tumor-bearing mice with the JAK1/2 inhibitor ruxolitinib [35] and quantified SIINFEKL presentation at the tumor cell surface. Ruxolitinib reduced the frequency of SIINFEKL⁺ tumor cells compared with vehicle-treated Ngfr-KO tumors (**Fig. 5d, e**), indicating that JAK activity contributes to the antigen-presentation phenotype observed upon NGFR loss *in vivo*. We therefore tested whether JAK signaling was also required for the immune phenotype induced by NGFR loss by treating MOC2 GL CTL and Ngfr-KO tumor-bearing mice with ruxolitinib [35].

Notably, ruxolitinib restored the growth of Ngfr-KO tumors, while having a more limited effect on CTL tumors (**Fig. 5f, g).** In addition, JAK inhibition reduced the accumulation of effector CD8⁺ T cells in Ngfr-KO tumors to levels comparable to those observed in CTL tumors **(Fig. 5h, i**). This phenotype indicates that the growth impairment of Ngfr-KO tumors depends on a JAK-associated antigen-presentation program that supports effective CD8⁺ T-cell immunity *in vivo*.

### NGFR blockade reverses anti-PD1 resistance in MOC2-derived tumors

Alterations in antigen presentation constitute a conserved mechanism of resistance to ICB in cancer [15, 36–41]. Since our data indicates that NGFR blockade enhances antigen presentation in tumor cells and promotes effector CD8^+^ T cells expansion, we next evaluated whether this NGFR-associated immune remodeling could influence response to anti-PD1 therapy. To evaluate the implications of this NGFR-associated phenotype in ICB responses, we treated MOC2 GL CTL or *Ngfr*-KO tumor-bearing mice with anti-PD1 or isotype control IgG every three days starting on day 6 post-implantation and continuing until the experimental endpoint. In line with previous data in the MOC2 model [31], CTL tumors did not respond to anti-PD1 treatment and continued to grow (**Fig. 6a, left panels; Fig. 6b, Extended Data Fig. 5a, b**). In contrast, Ngfr-KO tumors displayed a marked reduction in size following anti-PD1 treatment (**Fig. 6a, right panels; Fig. 6b, Extended Data Fig. 5c, d**). To evaluate response patterns, we quantified the change in tumor size at endpoint relative to the treatment start (D6). All CTL tumors were classified as non-responders, while 8 of 10 Ngfr-KO tumors exhibited a robust response with >75% reduction in tumor size (**Fig. 6c**). Notably, 6 of 10 Ngfr-KO tumors showed a complete response, with total tumor clearance (100% ± 1%) (**Fig. 6c**), indicating marked sensitization to ICB following NGFR depletion. Consistently, bulk RNA-seq comparing CTL and Ngfr-KO tumors revealed broad upregulation of immune-activated and APM-related pathways in Ngfr-KO tumors (**Extended Data Fig. 5e**), including IFN-γ response (Hallmark), MHC class I protein binding (GO:0042288), T cell–mediated cytotoxicity (GO:0001913) and antigen processing and presentation of peptide antigen (GO:0048002) (**Extended Data Fig. 5f–i**).

**Fig. 6.**
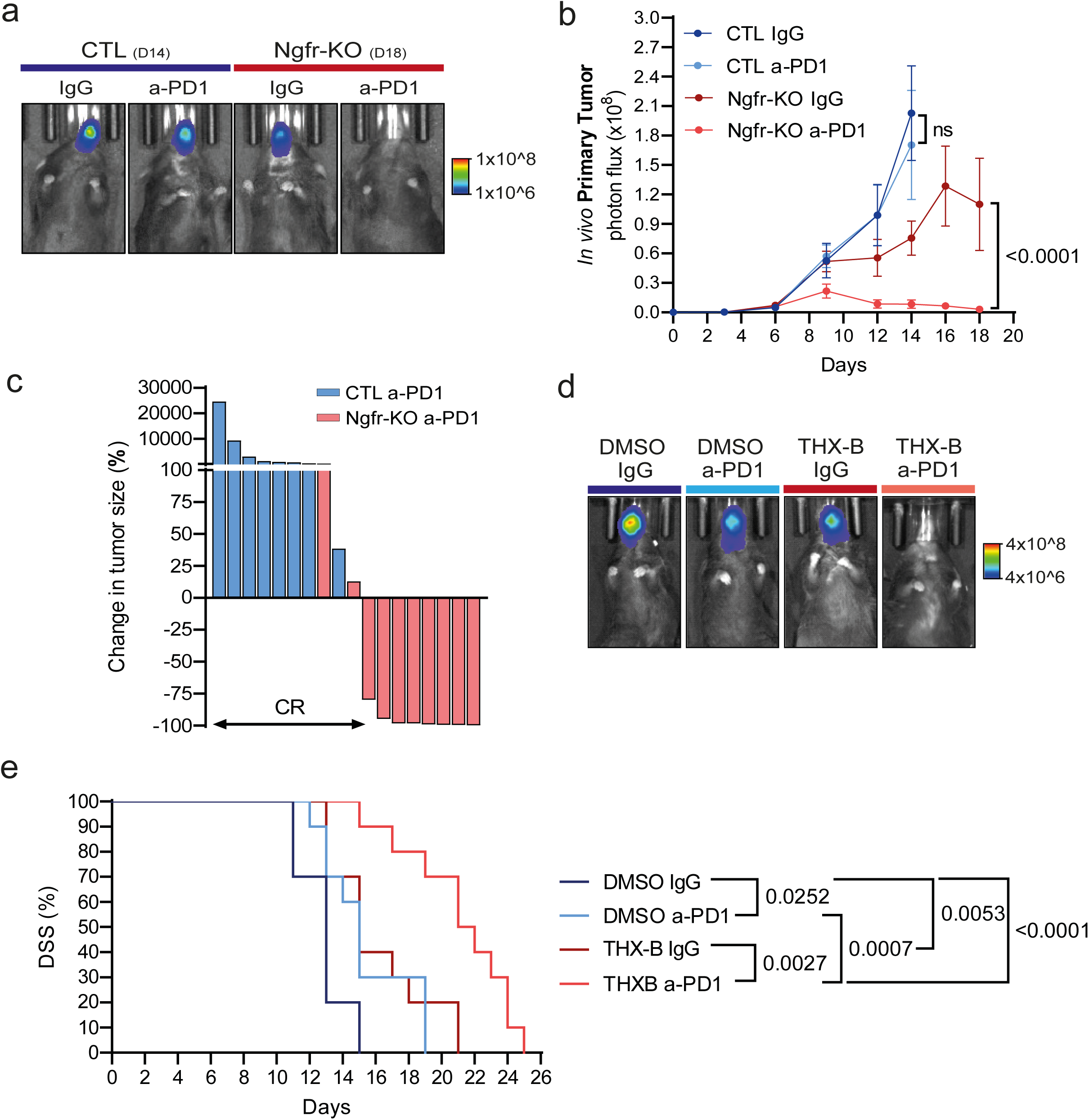
| Targeting NGFR reverts anti-PD1 resistance in MOC2 model. a, Representative endpoint BLI images from IgG- or anti-PD1-treated groups, showing CTL tumors at D14 (left) and Ngfr-KO tumors at D18 (right). b, Longitudinal tumor growth curves based on BLI, presented as mean ± SEM. Statistical comparisons were performed at each time point using two-way ANOVA. P values shown reflect comparisons at endpoint between CTL IgG and anti-PD1 at D14 and between Ngfr-KO IgG and anti-PD1 at D18. Sample sizes: CTL IgG (n = 10), CTL anti-PD1 (n = 9), Ngfr-KO IgG (n = 10), Ngfr-KO anti-PD1 (n = 10), from one experiment. c, Waterfall plot showing individual responses to anti-PD1 treatment, calculated as percent change in tumor size at endpoint relative to tumor size at treatment initiation (D6). Complete response was defined as a reduction of −100% ± 1%. Each line represents one mouse. Sample sizes: CTL anti-PD1 (n = 8; one mouse excluded due to unreliable measurement on D6), Ngfr-KO anti-PD1 (n = 10). d, Representative BLI images taken at D11 post-injection from MOC2 GL tumor-bearing mice treated with 0.05% DMSO or THX-B in combination with IgG or anti-PD1 (DMSO + IgG, DMSO + anti-PD1, THX-B + IgG and THX-B + anti-PD1). e, Kaplan– Meier survival plot representing disease-specific survival (DSS) of the same four treatment groups. The P values were calculated using a log-rank test. Sample sizes: n = 10 per group, pooled from two independent experiments.

Building on these results in the MOC2 GL Ngfr-KO model in combination with ICB, we next moved toward a more clinically relevant setting. We therefore evaluated tumor growth *in vivo* in mice orthotopically implanted with MOC2 GL cells and treated with THX-B or vehicle (0.05% DMSO) in combination with anti-PD1 or isotype IgG. To simulate a realistic therapeutic setting, we defined an individual endpoint for each mouse, based on signs of excessive tumor burden and/or weight loss, allowing us to assess disease-specific survival (DSS). Notably, the combination of THX-B with anti-PD1 led to a marked reduction in tumor growth (**Fig. 6d, Extended Data Fig. 5j–m),** which translated into a substantial improvement in survival, outperforming not only the control group (DMSO + IgG), but also each monotherapy alone (THX-B or anti-PD1) **(Fig. 6e**).

### NGFR is linked to tumor progression and impaired immune communication in melanoma

To explore whether the immunoregulatory role of NGFR extends beyond HNSCC, we analyzed its function in the immunocompetent Yumm1.1 melanoma model, which represents an immunotherapy-resistant melanoma system with intrinsic resistance to immune checkpoint blockade [17, 42, 43]. Yumm1.1 CTL and Ngfr-KO cells were injected subcutaneously into C57BL/6 mice, and tumor growth was monitored longitudinally until endpoint. Efficient NGFR deletion was confirmed by immunoblotting (**Extended Data Fig. 6a).** We confirmed that genetic ablation of Ngfr impaired primary tumor growth *in vivo* **(Extended Data Fig. 6b–d**).

Flow cytometry analysis of Yumm1.1 CTL and Ngfr-KO tumors revealed a significant increase in the effector CD8⁺ T-cell fraction compared with CTL tumors (**Extended Data Fig. 6e**). This finding is consistent with recent melanoma data showing that pharmacological inhibition of NGFR with THX-B increased intratumoral CD8⁺ T-cell infiltration [17].

### NGFR-linked antigen-presentation programs associate with local immune exclusion and clinical outcomes in HNSCC and melanoma patients

To assess the clinical relevance of NGFR in human disease, we performed a multi-level analysis of patient samples and datasets. This included: (i) evaluation of NGFR expression in human HNSCC tissues by immunohistochemistry; (ii) high-plex spatial profiling of HNSCC tumors to define the relationship between NGFR-positive malignant regions and the local immune microenvironment; and (iii) interrogation of human HNSCC and melanoma cohorts to determine whether an NGFR-regulated antigen-presentation signature is associated with clinical outcome and response to immune checkpoint blockade.

Analysis of NGFR expression in HNSCC tumors showed a predominant membranous staining in tumor cells with two expression patterns. In some cases, NGFR expression was restricted to a limited number of tumor cells in peripheral regions of the tumor in direct contact with the stroma, a pattern we termed peripheral (**Fig. 7a**). In contrast, other tumors showed a diffuse NGFR pattern, characterized by a higher proportion of NGFR⁺ tumor cells and stronger staining intensity throughout the tumor mass (**Fig. 7b**). Notably, the diffuse pattern of NGFR expression was associated with worse overall survival in cases with carcinoma in the hypopharynx (**Fig. 7c**). We also evaluated the association of NGFR with prognostic markers and metastatic disease. Diffuse NGFR pattern positively correlated with poor prognostic markers such as p-SRC [44–46], NANOG [47, 48] and β-nuclear catenin [49] (**Extended Data Fig. 7a**) and showed a negative correlation with good prognostic markers such as p-pS6 and p21 [50] (**Extended Data Fig. 7b**). Consistently, we observed an increased incidence of metastases in patients with NGFR diffuse pattern (**Fig. 7d** and **Extended Data Fig. 7c, d**).

**Fig. 7.**
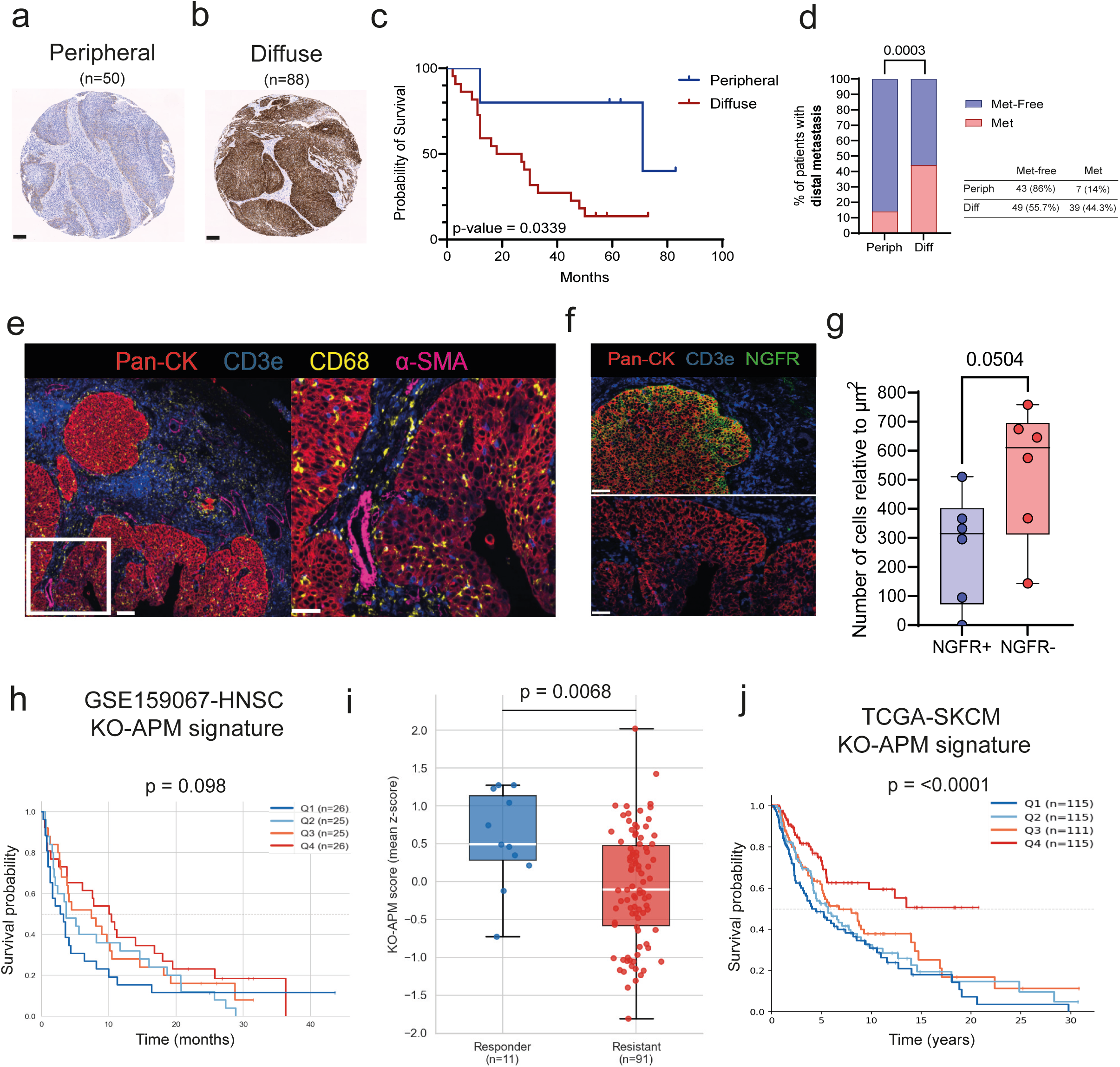
| NGFR identifies an aggressive and immunoevasive phenotype in human HNSCC patients. a, Representative image of one case of the TMA with peripheral NGFR expression (n = 50). b, Representative image of one case of the TMA with diffuse NGFR expression (n = 88). Scale bars, a–b: 100 µm. c, Kaplan–Meier curve showing overall survival of HPV-negative patients with hypopharyngeal cancer, stratified according to NGFR expression pattern. The P value was calculated using the log-rank test. d, Contingency plot showing the association between NGFR expression patterns and the presence of distant metastasis. A summary of the cases and percentages is shown below the plot. P value was calculated using Fisher’s exact test, comparing the total number of patients in each group. e, Representative high-plex images from one HNSCC patient. PanCK (red), CD3e (blue), CD68 (yellow) and α-SMA (pink) are shown. Scale bars: 100 µm (left) and 50 µm (right). f, Tumor regions from the same patient showing PanCK⁺ tumor areas that are NGFR⁺ (top) or NGFR⁻ (bottom). PanCK (red), CD3e (blue) and NGFR (green) are shown. Scale bars: 50 µm. All images were acquired with the MACSima™ system. g, Quantification of CD3⁺ T-cell infiltration in NGFR⁺ and NGFR⁻ tumor regions, normalized to the corresponding tumor area. Data are shown for n = 6 patients from two independent experiments. P value was calculated using a paired two-sided t-test, pairing values from the same tumors. h, Kaplan–Meier overall survival analysis of HNSCC patients treated with immune checkpoint blockade (ICB), stratified by the NGFR-KO–derived antigen-presentation signature (by quartiles). The P value was calculated using the log-rank test. i, Comparison of NGFR-KO antigen-presentation signature scores between responder and resistant patients in the ICB-treated HNSCC cohort. P value was calculated using a two-sided Wilcoxon rank-sum test. j, Kaplan– Meier overall survival analysis of melanoma patients from TCGA stratified by the NGFR-KO–derived antigen-presentation signature (by quartiles). The P value was calculated using the log-rank test.

To determine whether the adverse clinical outcome associated with the diffuse NGFR pattern reflects impaired antigen presentation and immune evasion, we selected individual cases exhibiting this expression pattern and characterized them by high-plex immunofluorescence. We characterized tumor intrinsic markers: Ki67, PD-L1 and HLA-DR to measure proliferative state, immune inhibitory ligand expression and tumor cell antigen-presenting capacity, respectively (**Extended Data Fig. 7f–h**) [51–53]. Importantly, we observed that PanCK⁺NGFR⁺ tumor cells exhibited lower HLA-DR expression compared to PanCK⁺NGFR⁻ cells, while no differences were observed in PD-L1 or Ki67 (**Extended Data Fig. 7e**). Additionally, we characterized several markers of the TME to evaluate NGFR-mediated immunosuppression (**Fig. 7e**). Importantly, NGFR⁺ regions showed a trend toward reduced infiltration of CD3⁺ T cells (**Fig. 7f, g**), while no differences were detected in macrophages (**Extended Data Fig. 7i**) or CD11b⁺ immune cells (**Extended Data Fig. 7j**). Together, these findings indicate that the diffuse NGFR pattern in human HNSCC is associated with reduced antigen-presenting capacity in tumor cells and selective immune exclusion within NGFR-positive tumor regions, implicating NGFR as a mediator of local immune evasion in the clinical setting.

To assess the translational relevance of the NGFR-regulated antigen-presentation machinery (APM) program identified *in vivo*, we derived a signature from genes upregulated in MOC2 Ngfr-KO tumor cells and evaluated its prognostic and predictive value across human HNSCC and melanoma cohorts. In the TCGA HNSCC cohort, high APM signature scores were not significantly associated with overall survival in the overall treatment-unselected population (**Extended Data Fig. 7k**), suggesting that this program is not a general prognostic marker in HNSCC. We therefore evaluated its relevance in the context of immune checkpoint blockade. In an independent ICB-treated HNSCC cohort, high APM signature scores showed a non-significant trend towards improved overall survival (**Fig. 7h**; p = 0.098), and responders showed significantly higher signature scores than non-responders (**Fig. 7i**). Extending this analysis to melanoma, we found that high APM signature scores were also associated with improved overall survival in TCGA melanoma (**Fig. 7j**), supporting the broader clinical relevance of this NGFR-linked antigen-presentation signature across tumor types.

## Discussion

NGFR has gained increasing recognition as both a marker and functional regulator of malignant plasticity, serving as a link between stem-like tumor states and metastatic progression as well as therapeutic resistance. In the context of melanoma, NGFR^+^ cancer cells exhibit an enhanced capacity to initiate tumor growth and metastasize[2, 54]. Additionally, NGFR marks EMT-associated stem cell states [3, 4] that are critical in resistance to therapies, such as chemotherapy [55], targeted therapies [56–59] and immunotherapies [14, 16, 17, 60].

In HNSCC, NGFR identifies a subpopulation characterized by tumor stem cell properties, enhanced metastatic potential, and a poorer prognosis [8, 10–12]. The identification of NGFR-positive stem-like populations in breast, colon, and esophageal cancers further implies that NGFR delineates a conserved plasticity program across various tumor types [5–7]. Interestingly, evidence across tumor types suggests that the subcellular localization and spatial pattern of NGFR, rather than its expression alone, can serve as a specific marker of tumor aggressiveness. In melanoma, NGFR^hi^ cells preferentially localize at the invasive front of metastatic tumors, where they exhibit vasculogenic mimicry and high PD-L1 expression in close proximity to immune cells, linking NGFR to invasion and immune evasion [16, 17]. However, a shift toward a diffuse intratumoral NGFR pattern is observed in more advanced disease contexts, including HNSCC, where it correlates with poor prognosis and metastatic burden, and in melanoma lung metastases, where NGFR expression becomes broadly distributed across metastatic lesions [16]. Similarly, the shift from restricted, myoepithelial NGFR localization in normal breast to aberrant expression in aggressive subtypes supports this concept [61]. Collectively, these studies suggest that NGFR serves not merely as a lineage or tumor-type-specific marker but as a broader indicator of aggressive, therapy-resistant tumor cell states, closely linked to its spatial redistribution into diffuse or invasion-associated areas in constant communication with immune cells. In this study, we expand upon this concept by demonstrating that NGFR also influences tumor–immune interactions, limiting antigen presentation and constraining CD8⁺ T-cell-mediated antitumor immunity.

Our findings place NGFR at the interface between malignant plasticity and tumor immune evasion, showing that this receptor regulates a tumor-cell state that restrains antigen presentation, limits CD8⁺ T cell-mediated adaptive immunity and influences the response to immune checkpoint blockade. This positions NGFR as a therapeutically actionable regulator of tumor immune resistance whose pharmacological modulation may enhance ICB responses. We found that *Ngfr* deletion shifts malignant cells from EMT/invasive states toward more epithelial and proliferative phenotypes, in line with previous HNSCC studies linking NGFR to EMT-like programs driven by factors such as ESM1 and SNAI2/Slug [11, 12, 62, 63]. Importantly, this apparent increase in epithelial/proliferative tumor-cell states occurs in tumors with reduced growth, indicating that NGFR loss uncouples intrinsic proliferative features from overall tumor progression.

Antigen presentation through MHC-I is a key requirement for effective antitumor immunity and clinical response to ICB [64, 65]. Defects in this pathway, including β2M mutations [36–38] and transcriptional silencing of MHC genes [15, 39, 40], are established mechanisms of immune escape and therapeutic resistance. Our results place NGFR within this axis by showing that *Ngfr* deletion triggers an antigen-processing and presentation program, encompassing MHC-I, MHC-II, TAP and immunoproteasome components. Importantly, we found that this is not merely transcriptional since Ngfr-KO tumor cells display increased surface presentation of the SIINFEKL peptide and improved recognition by tumor-specific CD8⁺ T cells. These data align with melanoma studies linking NGFR to reduced MHC-I expression, immune cell exclusion and resistance to T cell-mediated cytotoxicity [14, 16, 17].

The IFN-γ–JAK/STAT–IRF1 axis is a central pathway through which tumor cells become visible to T cells by inducing MHC-I/MHC-II expression and antigen-processing machinery components [65–71]. In OSCC/HNSCC, however, IFN-γ signaling appears to exert context-dependent effects on tumor immunity. On the one hand, IFN-γ can enhance tumor immune recognition by activating antigen-presentation programs in malignant cells. In line with this, recent single-cell analyses in HNSCC identified a malignant-cell IFN/MHC-II program associated with response to pembrolizumab, defining a clinically relevant immunogenic tumor-cell state linked to ICB sensitivity [72]. Similarly, OSCC studies have shown that IFN-γ can restore components of the antigen-processing machinery, including TAP1 and tapasin, and enhance antitumor immune responses [73], as well as induce MHC-II expression in head and neck cancer cells [74]. On the other hand, IFN-γ may also promote adaptive immune resistance, as illustrated by its ability to induce PD-L1 expression in oral squamous carcinoma cells through PKD-dependent signaling [75]. Thus, the functional outcome of IFN-γ signaling depends on the regulatory state of the tumor. Our data fit within this framework and adds a mechanistic layer by identifying NGFR as a negative regulator of the immunogenic arm of IFN-γ signaling. NGFR loss releases an IFN-γ/JAK-associated antigen-presentation program, increasing APM gene expression, tumor antigen presentation, effector CD8⁺ T-cell accumulation and immune-mediated tumor control. These findings suggest that NGFR acts as a brake on the IFN/MHC/APM program associated with favorable immune microenvironments and response to checkpoint blockade.

A key consequence of Ngfr loss is the conversion of poor immunogenic tumors into lesions capable of supporting productive CD8⁺ T-cell responses. In our model, Ngfr deletion selectively reshaped the CD8⁺ compartment toward an effector cytotoxic state, supporting a direct link between NGFR-dependent antigen presentation and adaptive immune control in HNSCC. This is consistent with melanoma studies showing that NGFR promotes resistance to T cell-mediated killing[14] and that NGFR inhibition improves PD-1/PD-L1 blockade efficacy [17], that IFN-γ-inducible CD271/NGFR suppresses melanoma antigen expression and limits T-cell activation [76], and that NGFR^high^/PD-L1^high^ tumor states are in close proximity to immune cells during acquired resistance to immunotherapy[16]. Our human analyses indicate that NGFR is associated with clinically relevant immune-evasive tumor states in both HNSCC and melanoma. In HNSCC, diffuse NGFR expression identified aggressive tumors with increased metastatic incidence and poorer clinical outcome. In line with this, an NGFR-regulated antigen-presentation machinery signature was enriched in HNSCC patients responding to immune checkpoint blockade. Thus, our data expand the role of NGFR in HNSCC beyond its previously described association with disease progression and macrophage-mediated remodeling of the tumor immune microenvironment[77], revealing a link between NGFR-positive tumor states, defective antigen-presentation programs and reduced immune visibility. In melanoma, previous studies have shown that tumor-intrinsic NGFR signatures predict anti–PD-1 resistance and that NGFR^high^ tumor cells are associated with immune exclusion[14]. Consistent with this framework, our NGFR-regulated antigen-presentation machinery signature was associated with improved survival in melanoma patients, supporting the clinical relevance of this NGFR-linked immune-visibility program across tumor types.

From a therapeutic perspective, NGFR/CD271 is an actionable vulnerability in aggressive and therapy-resistant tumor-cell states. Initial studies in HNSCC established CD271 as a functional and targetable marker of tumor-initiating cells, showing that CD271 inhibition reduces ERK signaling and impairs tumor initiation *in vivo*[9]. In melanoma, antibody-based strategies have been explored, including combined CD47 blockade and CD271 targeting in patient-derived xenografts, which suppresses metastatic dissemination[78], as well as humanized anti-CD271 antibodies capable of depleting CD271⁺ cancer stem-like cells through antibody-dependent cytotoxicity[79]. Other approaches have activated the CD271 intracellular death domain in combination with chemotherapy or targeted therapy to inhibit melanoma progression[80]. In parallel, indirect approaches that reduce NGFR expression or NGFR-associated phenotype switching, such as HSP90 inhibition, MNK1/2–eIF4E blockade or metabolic rewiring with ranolazine, have been shown to enhance antitumor immune responses and improve immunotherapy sensitivity [14, 81, 82]. In addition, p75NTR/NGFR can be directly targeted with small molecules [83], blocking melanoma migration and lung invasion[84]. Our work showed that pharmacological NGFR inhibition using the small molecule THX-B reduces melanoma local and distal metastasis, increasing intratumoral CD8⁺ T-cell infiltration and restores sensitivity to PD-1/PD-L1 blockade in immunotherapy-resistant models [17]. Our data support that NGFR targeting restrains primary tumor growth and spontaneous metastasis, promoting effector CD8⁺ T-cell responses and sensitizes otherwise resistant tumors to anti-PD1 therapy. This study supports NGFR inhibition as a strategy to simultaneously target aggressive NGFR-dependent tumor states and restore tumor immune visibility, thereby improving response to immune checkpoint blockade.

## Methods

### Cell lines and culture

The parental murine model of HNSCC MOC2 [30] was provided by Dr. Aznar-Benitah (Institute for Research in Biomedicine, Spain). The murine melanoma cell line Yumm1.1 [42] was kindly provided by Dr. Soengas (Spanish National Cancer Research Centre, Spain). MOC2 cells were cultured in a 2:1 mixture of Iscove’s Modified Dulbecco’s Medium (IMDM) (SH30228.02, Fisher Scientific) and Ham’s F12 Nutrient Mixture (F12) (SH30026.01, Fisher Scientific), supplemented with 10% fetal bovine serum (FBS, Thermo Fisher Scientific), 40 µg/mL hydrocortisone (H0135-1MG, Sigma-Aldrich) and 5 µg/mL insulin (I6634-50MG, Sigma-Aldrich). Yumm1.1 cells were cultured in a Ham’s F12 and Dulbecco’s Modified Eagle Medium (F12/DMEM) mixture supplemented with MEM Non-Essential Amino Acids Solution (#11140-050, Gibco) and 10% FBS. Additionally, gentamicin (Sigma-Aldrich; 200 µg/mL), glutamine (Sigma-Aldrich; 2 mM) and sodium pyruvate (Sigma-Aldrich; 10 mM) were added to all cell lines. Cells were maintained at 37 °C in a humidified atmosphere with 5% CO₂ and subcultured using trypsin-ethylenediaminetetraacetic acid 0.05% (EDTA) (Thermo Fisher Scientific).

### Genetically modified cell lines

MOC2 cells were transduced with a green fluorescent protein (GFP)–luciferase (Luc) reporter cassette (luciferase–IRES–EGFP in pMSCV luc gfp). Knockout (KO) cell lines for MOC2 and Yumm1.1 were generated using the lentiCRISPRv2 system (Addgene plasmid #52961) with single-guide RNAs (sgRNAs) targeting Ngfr (lentiCRISPRv2-Ngfr) or Rosa26 (lentiCRISPRv2-Rosa26); Rosa26-targeted cells were used as CRISPR control (CTL). For the ovalbumin (OVA)–mCherry model, both MOC2 GFP–Luc CTL and Ngfr-KO cell lines were transduced with the lentiviral vector pLEX-OVAIC (pLEX-OVA–IRES–mCherry). Lentiviruses were prepared as previously described [85]. Briefly, 9 μg of the lentiviral plasmid, purified with the EndoFree Plasmid Maxi Kit (QIAGEN, USA), was transfected into HEK293T cells together with 3 μg pMD2.G (Addgene plasmid #12259), 5 μg psPAX2 (Addgene plasmid #12260) and 62.5 μl Lipofectamine™ 2000 (Life Technologies). At 48 h, the supernatant containing the lentiviruses was recovered and filtered through a 0.45 μm low protein binding filter (Millipore). Cells (100,000 per well) were transduced in suspension with the lentiviral supernatant at a low multiplicity of infection (MOI) in the presence of 8 µg/ml polybrene. GFP-positive cells were expanded and sorted on a BD Influx cell sorter. Cells transduced with lentiCRISPRv2 constructs were selected with 1 µg/ml puromycin (InvivoGen). NGFR deletion was confirmed by Western blot, and NGFR-KO cells were further enriched by sorting for NGFR-negative cells on a BD Influx cytometer. Cells transduced with OVA– IRES–mCherry (OVA–mCherry) were selected with 1 µg/ml hygromycin B and subsequently sorted for mCherry expression on a BD Influx™ cell sorter. The efficiency of OVA–mCherry expression was confirmed by fluorescence microscopy. Additional information on the constructs is summarized in **Supplementary Table 1**.

### Animal studies

All the experimentation involving mice were approved by the Comunidad Autónoma de Madrid (PROEX029.5/22 and PROEX82.7/23). Tumor burden did not exceed a maximum volume of 1 cm3 or a BLI signal of 1.5 x 10⁹ photons in primary tumor growth or metastatic spread studies. The experiments were performed in accordance with the guidelines for Ethical Conduct in the Care and Use of Animals as stated in The International Guiding Principles for Biomedical Research involving Animals, developed by the Council for International Organizations of Medical Sciences.

C57BL/6JOlaHsd mice were bred at the CNIO mouse facility. Batf3-/-mice (Batf3tm1Kmm) were originally generated by K. M. Murphy (Washington University School of Medicine in St. Louis). OT-I mice (C57BL/6JOlaHsd-PtprcaRag1ko/koTg(TcraTcrb)1100Mjb/J) were originally generated by M. J. Bevan (University of Washington Medical Center). OT-I mice were maintained on a CD45.1 background for adoptive transfer studies. Eight- to 12-week-old mice were used for all experiments.

### Orthotopic tumor implantation

For orthotopic implantation, 10^4^ MOC2 cells were injected into the tongue of each mouse. The cells were resuspended in serum-free IMDM/F12 medium. Mice were previously anesthetized intraperitoneally (i.p.) with ketamine (100 mg/kg) and medetomidine (1 mg/kg). Atipamezole (1 mg/kg) was administered i.p. as an antagonist to reverse sedation. Analgesic buprenorphine (0.1 mg/kg) was administered subcutaneously 15 min before the procedure, and then every 12 hours for the next 48 hours.

*In vivo* tumor growth was monitored two to three times per week by bioluminescence imaging (BLI). D-luciferin (150 mg/kg) was administered i.p., after which mice were anesthetized with 4% isoflurane and imaged in an *in vivo* imaging system (IVIS) system. The total luminescence of each region of interest (ROI) was quantified using Living Image software. ROIs were manually defined on the bioluminescent signal within the oral cavity. Animals were maintained until day 13 or 14, which was set as the experimental endpoint, determined from the onset weight loss in control mice. Using this same temporal scheme as a common endpoint for all groups, spontaneous metastasis was assessed by BLI. To this end, *ex vivo* imaging of the cervical tumor-draining lymph nodes (tdLNs) and lungs was performed after D-luciferin injection.

In the orthotopic experiments aimed at assessing survival, tumor growth was monitored in the same way by BLI. However, in this case, the endpoint was defined individually for each animal. The predefined criterion established was a weight loss of more than 10 % and/or a bioluminescence signal greater than 1.5 x 10⁹ photons/second in the total luminescence measurement (total flux).

### Subcutaneous tumor implantation

For subcutaneous injection of tumor cells, mice were anesthetized with 4% isoflurane. 10^6^ Yumm1.1 CTL or Ngfr-KO cells were resuspended in a 1:1 mixture of Matrigel and phosphate-buffered saline (PBS) and injected subcutaneously into the flanks of C57BL/6 mice. To evaluate tumor growth, the width and length of each tumor were measured twice a week using a digital caliper. Tumor volume was calculated using the formula V = a x *b*^2^ x 0.5236, where “a” represents the longest dimension and “b” the shortest. A common endpoint was established for all animals, defined as the time at which the CTL tumors reached a volume of 1 cm3.

### Tail vein tumor implantation

For experimental metastasis evaluation, tumor cells were injected into the lateral tail vein using a 26G needle. Prior to injection, the tail of each animal was exposed to a heat lamp for 5 min to promote vasodilation, and mice were immobilized in a dedicated restraining device during inoculation. A total of 1 × 10⁵ MOC2 GL CTL or Ngfr-KO cells were resuspended in serum-free medium and were injected per mouse. Metastasis was monitored *in vivo* once a week by BLI as described above. Prior to each acquisition, the thoracic region of the mice was shaved using an electric shaver to optimize signal detection. After four weeks, the experimental endpoint was reached, at which time the presence of lung metastases was assessed *ex vivo* by BLI.

### OT-I cells adoptive transfer

To assess antigen-specific CD8⁺ T cell responses *in vivo*, we performed adoptive transfer of OT-I cells into tumor-bearing mice. MOC2 GL OVA–mCherry CTL or Ngfr-KO cells were orthotopically implanted into the tongues of C57BL/6 (CD45.2) mice as described above. On day 4 after tumor implantation, spleens were collected from OT-I × CD45.1 donor mice, mechanically dissociated to obtain single-cell suspensions, and subjected to red blood cell lysis with ACK lysing buffer (Lonza) for 1 min at room temperature (RT), followed by washing and counting of splenocytes. CD8⁺ T cells were then enriched by positive selection using mouse CD8 (Ly-2) MicroBeads and LS MACS columns (Miltenyi Biotec, 130-117-044 and 130-042-401, respectively). Purified OT-I CD8⁺ T cells were resuspended in sterile PBS, and 10⁶ cells were injected intravenously (tail vein) into each tumor-bearing mouse on day 4 after tumor implantation.

### *In vivo* treatments

For treatment experiments with THX-B and/or anti-PD1, the following regimen was used. Six days after orthotopic injection of tumor cells, once a clearly visible tumor was detected by BLI in the oral cavity of all mice, treatment was initiated. THX-B was resuspended in a solution of PBS with 0.05% dimethyl sulfoxide (DMSO). THX-B was administered i.p. at a dose of 100 µg/mouse, or vehicle (0.05% DMSO in PBS) was injected as a control. For immunotherapy treatment, an anti-PD1 monoclonal antibody (InVivoPlus, clone RMP1-14, BioXCell #BE0146) administered i.p. at 200 µg/mouse was used. A rat immunoglobulin G (IgG)-2a isotype control (BioXCell #BE0090) was used as a control. Both THX-B and anti-PD1 treatments with their respective controls were administered every three days.

For CD8^+^ T cell depletion assays, mice were injected i.p. with an anti-CD8 monoclonal antibody at a dose of 200 µg/mouse (BioXCell #BE0223) or a control IgG isotype (BioXCell #BE0090) one day prior to tumor inoculation.

To evaluate the role of the JAK1/2-STAT1 pathway, mice were treated i.p. with ruxolitinib (INCB018424). Based on stability assays, the compound was freshly prepared every 5 days as a homogeneous suspension in sodium carboxymethyl cellulose (CMC-Na). Mice received one-daily administration at a dose of 900 µg/mouse. The suspension was maintained under agitation during administration and stored at 4°C.

### Tumor dissociation and cell suspension

Tumors were processed to obtain a single-cell suspension by mechanical and enzymatic digestion. Initially, tissues were mechanically disaggregated using scalpels and subsequently incubated at 37 °C in a shaking water bath for 20 min in 4 mL of an enzymatic mixture consisting of 0.8 mg/mL Dispase II (Roche), 0.2 mg/mL Collagenase P (Roche) and 0.1 mg/mL DNase I (Roche), dissolved in RPMI-1640 (Sigma). After the first digestion, the tumors were further disaggregated by pipetting and repeated passage through syringes. The material was allowed to stand at 4 °C for a few minutes to sediment the tissue debris, after which the supernatant was collected and transferred to a new tube containing fluorescence-activated cell sorting (FACS) buffer (PBS with 5 mM EDTA and 0.1 % bovine serum albumin (BSA), Sigma). The remaining sediment was again incubated with 4 mL of enzyme mixture for 10 min at 37 °C with shaking. After this second digestion, the tissue debris was disaggregated as before and combined with the initial supernatant. The resulting suspension was centrifuged at 300 × g for 4 minutes at 4°C. The cell pellet was resuspended in ACK red blood cell lysis buffer (Lonza) for 1 minute for erythrocyte lysis, diluted with 10 mL of FACS buffer, and centrifuged again under the same conditions. Finally, the pellet was resuspended in FACS buffer and filtered through 70 μm mesh filters (Corning) to remove cell aggregates.

### Flow cytometry

After obtaining a single-cell suspension from the tissues, cells were counted using a Countess 3 FL automatic cell counter (Invitrogen). For each experiment, the same number of cells per sample was stained, usually 10^6^ cells. First, cells were incubated for 15 min at 4 °C with anti-mouse CD16/CD32 Fc receptor blocking antibody (BD) diluted 1:50 in 100 μL of FACS buffer (PBS, 5 mM EDTA, 0.1% BSA (Sigma)), in 96-well round-bottom plates (Thermo Fisher Scientific). They were then washed with FACS buffer and centrifuged for 4 min at 300 g and 4 °C. Cell pellets were resuspended in the experiment-specific antibody mixture and incubated for 30 min at 4 °C in the dark. Subsequently, cells were washed with FACS buffer, centrifuged again and filtered using 50 μm CellTrics™ filters (Sysmex), transferring the suspension to 5 mL polystyrene round-bottom cytometry tubes (Corning). Data acquisition was performed on a BD LSRFortessa cytometer using FACSDiva v9.0 software (BD) or on a Cytek Aurora flow cytometer using SpectroFlo® software (Cytek Biosciences). Spectral compensation was performed with UltraComp eBeads™ (Invitrogen). Data was analyzed with FlowJo v10.10 software (Tree Star). Flow cytometry antibodies used are detailed in the **Supplementary Table 1**.

### Immunoblotting

Cells were lysed in 1× RIPA buffer (0.1% SDS, 0.5% sodium deoxycholate, 1% NP-40 (Igepal-CA-630), 150 mM NaCl, 50 mM Tris-HCl pH 8, and protease and phosphatase inhibitor cocktails (Sigma-Aldrich)) for 20 min on ice. Lysates were then centrifuged at 15,000 rpm for 20 min at 4 °C. Supernatants were used to quantify protein using bicinchoninic acid (BCA) assay (PierceTM BCA Protein Assay Kit; Thermo Scientific). Afterwards, 20 μg of protein in 1× Laemmli buffer were separated by sodium dodecyl sulfate–polyacrylamide gel electrophoresis (SDS–PAGE) and transferred to a nitrocellulose membrane (GE Healthcare). Membranes were blocked with 5% non-fat dry milk diluted in Tris-buffered saline with Tween 20 (TTBS). Membranes were then incubated with the primary antibodies overnight at 4°C. Subsequently, membranes were washed three times for 5 min with TTBS buffer and incubated with secondary antibodies for 1 hour at room temperature. Proteins were visualized using an enhanced chemiluminescence (ECL) detection system (GE Healthcare).

The primary antibodies used were the following: human NGFR (Abcam, ab52987; 1:1000), mouse NGFR (Monoclonal Antibody Core Unit, CNIO, NORI146C, 1:1000) and β-actin (Santa Cruz; sc-47778; 1:10000) as loading control. The Horseradish Peroxidase linked secondary antibodies used were the following: sheep anti-mouse IgG (NA931V, Lot15934109; 1:5000) and donkey anti-rabbit (NA934V, Lot 17212129; 1:5000).

### Single cell RNA sequencing

#### Experimental Design

MOC2 GFP-Luc CTL and Ngfr-KO cells were orthotopically injected into the oral cavity of C57BL/6J wild-type (WT) mice (n=10/group), following the conditions described above. Tumor growth was monitored by BLI and, on days 13 and 14 post-injection, the animals were sacrificed and the tumors removed from the oral cavity. On each experimental day, an independent cohort of animals was used, and tumors from 5 mice per group (CTL and Ngfr-KO) were pooled. The preparation of single-cell suspensions from tumors for flow cytometry was performed as described above.

Three different experimental conditions were used to obtain individual cells: (1) tumor cells, isolated by sorting for GFP⁺ (n = 2 per condition); (2) immune cells, isolated by sorting for CD45⁺ (n = 2 per condition); (3) enrichment of rare immune populations, in which CD45⁺ cells were sorted excluding neutrophils (Ly6G⁺, CD11b⁺) and macrophages (CD64⁺), with the aim of enriching lymphocytes and other less abundant myeloid populations (n = 1 per condition). The third enrichment condition was incorporated within the total CD45^+^ group for pooled analysis of immune cells. Additionally, a CD45^+^ healthy tongue was included as a control of the tumor-specific immune infiltration. A total of 2 x 10^5^ cells from each condition were sorted using the BD FACSAria Fusion Class II.

Single cell transcriptomic profiling was performed using 10x Genomics’ Chromium GEM-X Single Cell Gene Expression (3’ v4) technology. This technology allows the generation of Gel Beads-in-emulsion (GEMs), where each cell is encapsulated together with an oligonucleotide-coated bead containing a cellular barcode (10x Barcode), a Unique Molecular Identifier (UMI) and a poly(dT) sequence to capture mRNA. Samples were processed on the Chromium X instrument following the standard protocol for GEM-X 3’ v4. Reverse transcription and incorporation of cellular barcodes were performed within the GEMs, allowing the cellular identity of each transcript to be maintained. The generated libraries were amplified and prepared for sequencing short-read Illumina platforms.

#### Preprocessing

Alignment of FASTQ files and generation of gene expression matrices were performed using the standard Cell Ranger pipeline (10x Genomics), using a mouse genomic reference modified to include the GFP sequence. CellBender (v2.0) was then applied for environmental RNA correction and accurate cell identification, discarding cells with a cell probability of less than 0.8, thus minimizing noise from empty or contaminated droplets. Doublet detection was carried out using Scrublet, a tool based on transcriptomic profiling simulation. As an additional quality control step, filters were applied with Scanpy, eliminating cells with a mitochondrial content higher than 10% (indicative of cell damage or apoptosis) and those with less than 500 or more than 7,500 genes detected (which could correspond to empty droplets or undetected doublets). Once filtering was completed, each sample was preprocessed individually in Scanpy, including: 1) normalization for sequencing depth; 2) selection of highly variable genes (HVGs), considering approximately 10% of the genes detected per sample; 3) dimensionality reduction by PCA (50 principal components); 4) unsupervised clustering using the Leiden algorithm, testing different resolutions (0.1, 0.2 and 0.3).

Initial cell annotation was performed per sample, using a panel of markers defined by Inés Sentís (CNAG) in the same model, in collaboration with Salvador Aznar (IRB Barcelona). In tumor samples, clusters containing an appreciable proportion of GFP+ cells were classified as malignant, except for small peripheral clusters with a low proportion of GFP, which were annotated following the same criteria as non-tumor samples. Subsequently, data from all samples were integrated using Harmony, followed by global reclustering. After integration, small clusters with high doublet scores were identified. Therefore, cells with doublet score > 0.15 were removed and both integration and reclustering were repeated. Finally, a comprehensive re-annotation was performed at the cell and cluster level, combining automatic classification based on previously defined markers with refined manual annotation.

#### Functional analysis of malignant cells

For the analysis of tumor cells, we selected those annotated as “Malignant cells” derived from the GFP^+^ sorting that had passed the previous quality control filters. Using Scanpy, the data were first scaled (max value = 10), followed by PCA with 20 components. A two-dimensional representation was constructed using Uniform Manifold Approximation and Projection (UMAP). The Leiden clustering algorithm was applied with a resolution of 0.3, which allowed the identification of distinct cell groups while maintaining a biologically interpretable granularity. Subsequently, differential expression analysis was performed among the Leiden clusters using the Wilcoxon test. Clusters identified as contaminants were excluded because of high expression of immune or stromal genes, resulting in eight tumor clusters. Each of the clusters was manually annotated based on differentially expressed genes and supporting literature. Tables of cell counts per genotype (CTL and Ngfr-KO) as well as their respective proportions were calculated. To compare the distribution of tumor subtypes between the two conditions (CTL and Ngfr-KO), we calculated fold change, absolute percent change and increase in cell number.

To assess transcriptomic pathways deregulated according to genotype in tumor cells, Gene Set Enrichment Analysis (GSEA) was employed. A differential expression analysis was first performed between all malignant cells in the CTL and Ngfr-KO groups, and the results were filtered to retain only coding murine genes (Mus_musculus.GRCm39.113). From the ranked list of coding genes, enrichment analysis was performed using the GSEApy tool using a prerank approach. The analysis was applied separately to three murine databases: Hallmarks (h.all.v2024.1), Reactome (c2.cp.reactome.v2024.1) and Gene Ontology (c5.go.bp.v2024.1). The parameters used included 1,000 permutations and a range of 5-5,000 genes per pathway.

#### Functional analysis of immune cells

After data preprocessing and integration, CD45^+^ immune cells were grouped into distinct clusters based on the expression of classical marker genes. Twelve clearly defined immune populations were identified, both by their location in the reprojected UMAP (reUMAP) and by the differential expression of characteristic genes. These populations were annotated as: B cells, Dendritic cells, Innate Lymphoid Cells (ILCs), Macrophages, Mast cells, Monocytes, NK cells, Neutrophils, Progenitor ILCs, Senescent neutrophils, T cells and monocyte-derived Dendritic Cells (moDCs). Tables of absolute counts and relative proportions per genotype (CTL and Ngfr-KO) were generated. To evaluate differences between conditions, metrics such as fold change (FC) and absolute change in percentage of cells were calculated for each immune population.

Subsequently, a more detailed analysis of the T cell compartment was performed. Cells previously annotated as T cells were extracted, and a new UMAP projection was generated after independent preprocessing, including PCA (20 components) and clustering with the Leiden algorithm (resolution = 0.3). One of the clusters was excluded as it was considered contaminating, since it lacked expression of CD3e, an essential marker of T lymphocytes. A differential expression analysis was performed between the Leiden clusters using the Wilcoxon test, identifying the most characteristic genes of each group. Based on these genes and literature references, ten functional T-cell subtypes were defined and annotated as “CD4 helper”, “CD4 resting/memory”, “CD4 Tregs”, “CD4 Trm-like”, “CD8 cytotoxic”, “CD8 exhausted”, “CD8 naïve-like”, “IL-1 inflammatory”, “Proliferating” and “Th17-like”. Finally, the proportions of each subtype were compared between experimental conditions (CTL vs Ngfr-KO) by calculating the fold change and absolute percent change in their abundance.

### Bulk RNA sequencing

For bulk RNA-seq analysis, MOC2 GL CTL or Ngfr-KO tumors, treated or untreated with anti-PD1, were used to generate total RNA libraries that were sequenced on the Illumina NextSeq 2000 platform (P3 kit, 50 cycles), with 50 bp single-end reads, reverse strand-specific design, and a target of ∼50 million reads per sample, in two sequencing rounds per sample. The raw reads underwent preprocessing, alignment to the Mus musculus reference genome (GRCm39), and gene-level count assignment using a standard pipeline described in the preprocessing report. All samples passed quality control. A Python workflow was implemented. First, genes with a total count of zero were removed and then we extracted the set of coding genes, restricting the matrix to 18,418 coding genes. From this matrix, only untreated samples were selected. With these six samples and the coding genes, a differential expression model was fitted using pydeseq2 (DeseqDataSet with design ∼Condition), estimating size factors, dispersions, and log2FC, and applying the Wald test for the CTL vs. Ngfr-KO contrast. The results were filtered, retaining genes with baseMean ≥ 10, and those with padj < 0.05 and |log2FoldChange| > 0.5 were defined as differentially expressed.

### HNSCC TMA cohort

We analyzed a cohort generated at the Hospital Universitario Central de Asturias, Oviedo, Spain, consisting of tumors from 382 HNSCC patients [86, 87]. All patients presented a single primary tumor and did not receive prior treatment before surgery. HPV status was available for all patients. The tissue microarray (TMA) consisted of 1 mm tumor cores constructed following previously described methodologies [86, 87]. At the time of NGFR immunohistochemistry, the TMA comprised 379 patients across different anatomical locations: 249 from the oropharynx, 62 from the hypopharynx, and 68 from the larynx. After excluding HPV-positive cases (n = 10) and samples with insufficient tumor tissue, the final cohort included 330 evaluable patients, with 221 from the oropharynx, 56 from the hypopharynx, and 53 from the larynx. All patients had clinical follow-up data, including survival status (alive or deceased), follow-up duration in months, presence of nodal metastasis (N), presence of distant metastasis (M), and tumor recurrence. Pathological evaluation of NGFR staining identified positivity in 138 cases, with distinct membranous and cytoplasmic staining observed in tumor cells. The pathologist categorized NGFR expression into two patterns: a peripheral pattern, where NGFR expression was restricted to the outer region of the tumors in contact with the stroma, observed in 50 cases, and a diffuse pattern, where NGFR expression was widespread throughout the tumor, involving a larger proportion of tumor cells, observed in 88 cases.

This TMA cohort is characterized by immunohistochemistry for several prognostic markers, which have been evaluated and published [48–50]. We evaluated the associations of these markers with NGFR expression. Among the relevant markers associated with poor prognosis and malignancy are phosphorylated SRC (p-SRC), NANOG, and nuclear β-catenin, while markers associated with good prognosis include p21, p-pS6 (S235), and p-pS6 (S240).

### Immunohistochemistry

Tissues were harvested and fixed in 10% formalin (4% formaldehyde in solution) in segmented portions for 24 hours. Afterwards, tissues were transferred to 50% ethanol overnight before embedding into paraffin blocks. Samples were sectioned at 2.5 µm thickness, mounted on Superfrost® Plus slides, and dried ON. Immunohistochemical reactions were carried out using automated staining platforms (Discovery ULTRA, Ventana-Roche, or Autostainer Link 48, Dako). Antigen retrieval was performed, and endogenous peroxidase activity was blocked using 3% hydrogen peroxide.

Murine samples were incubated with an NGFR antibody (NGFR NORI146C Monoclonal Antibody Core Unit, CNIO). For detection of NGFR in the characterized patient cohorts, pre-sectioned slides were provided by Dr. Garcia-Pedrero and Dr. Alvarez-Fernandez and stained with a human NGFR antibody (Anti-p75 NGF Receptor antibody [NGFR5], Abcam; ab3125). Subsequently, they were incubated with the corresponding secondary antibodies (anti-rat (AI-5000-1.5, Vector) or anti-mouse (ab133469, Abcam) conjugated with horseradish peroxidase. The reaction was visualized using 3,3’-diaminobenzidine tetrahydrochloride (DAB) with Envision FLEX, Novolink polymer or OmniMap Discovery Ultra platforms, according to the manufacturers’ protocols. Nuclei were counterstained with Harris hematoxylin. Positive control sections were included in each staining. Representative images were acquired by brightfield microscopy.

### Immunofluorescence high-plex imaging

ROIs were selected from human HNSCC tumors previously characterized by diffuse NGFR expression. Selection was based on morphological quality of the tissue, homogeneity of staining and absence of artifacts. Each ROI was analyzed by cyclic immunofluorescence using the automated MACSima™ platform. Images were acquired at a resolution of 0.17 µm/pixel (20× objective, 0.45 numerical aperture) and stored in multichannel Open Microscopy Environment Tagged Image File Format (OME-TIFF) format. Nuclear segmentation was performed from the DAPI channel using Cellpose (Cyto3 model) via QuPath. To capture membranous and cytoplasmic markers, a morphological expansion of 3 pixels around the nucleus was applied. A labeled mask was generated per cell with unique identifiers that served as the basis for per cell quantification. Antibodies are summarized in **Supplementary Table 1**.

The analysis was performed using Python. Both the original multiplex images and nuclear segmentation masks generated from the DAPI channel were loaded. To identify tumor cells, PanCK⁺ masks were generated by applying a fixed intensity threshold (2,500). In the case of NGFR, due to inter-image variability, an adaptive threshold per ROI defined as: threshold = mean + score × standard deviation, where the score was manually adjusted for each image. In both cases, the masks underwent morphological postprocessing that included closing operations, removal of small objects (< 2,000 pixels) and gap filling (< 10,000-75,000 pixels), with the aim of obtaining clean, binarized masks. NGFR masks were further restricted to nuclear areas identified by DAPI, to avoid non-specific background signal. From the labeled DAPI masks individual cells were identified in each ROI. For each cell the following properties were quantified: area (in pixels and µmZ), centroid coordinates (X, Y), average intensity for each channel, and binary positivity status for CK and NGFR, by overlaying with their respective masks. Binary classification of markers was performed using adaptive thresholds per ROI, calculated as: threshold = mean_ROI + score × deviation_ROI, where the score (range 0-3) represents a user-defined degree of sensitivity for each marker. Subsequently, combinations of markers were defined to identify cell subpopulations of interest, and absolute counts and relative proportions per ROI were calculated. Finally, functional ratios were estimated within each subpopulation to assess phenotypic differences between cell groups.

### Statistics and Reproducibility

For the comparison of continuous values between two groups, an unpaired two-tailed t-test was used. When continuous values were compared between three or more groups, a one-way Analysis of Variance (ANOVA) with correction for multiple comparisons using Tukey’s test was applied. For the analysis of tumor growth between two or more groups over different time points, we used a two-way ANOVA with correction for multiple comparisons using Sidak’s test. Survival curves were analyzed using the Kaplan-Meier method, and statistical significance was assessed with the log-rank test (Mantel-Cox). To compare frequencies between groups (number of cases per condition), contingency analysis was used using the chi-square test, or Fisher’s exact test when the expected frequencies were low.

To identify the differentially expressed genes between different conditions in scRNA-seq data, the Wilcoxon rank-sum test was used. Frequency differences between genotype conditions (CTL and Ngfr-KO) at low categorical counts (<5) were evaluated with Fisher’s exact test, whereas for higher frequencies the chi-square test was used. The p-values were adjusted for false discovery rate (FDR) using the Benjamini-Hochberg method, considering as statistically significant and biologically relevant those results with adjusted p-value < 0.05 and an absolute percentage change greater than 5 %. Finally, assessment of the statistical significance of scRNA-seq-enriched pathways between CTL and Ngfr-KO malignant cells was performed by a permutation test on the Enrichment Score of the preranked GSEA analysis, followed by correction for FDR using the Benjamini-Hochberg method.

## Data availability

Single-cell and bulk RNA-sequencing data have been deposited in the NCBI Gene Expression Omnibus (GEO). Custom analysis scripts used in this study have been deposited on GitHub and will be made publicly available upon peer-reviewed publication.

## Acknowledgements

Dr Peinadós laboratory is supported by grants from Agencia Estatal de Investigación, Ministerio de Ciencia e Innovación / State Research Agencia Financiadora (AEI - Ministry of Science and Innovation: PID2020-118558RB-I00 (MICIU/AEI/10.13039/501100011033), RETOS SAF2017-82924-R (AEI/10.13039/501100011033/FEDER-UE), PDC2021-121102-I00 (MCIN/AEI/10.13039/501100011033, European Union “NextGenerationEU”/PRTR), PID2023-147503OB-I00 funded by MCIU/AEI /10.13039/501100011033 and by the European Union, ERDF “A way of making Europe”, Fundación Científica AECC (PRYCO223002PEIN, LABAE19027PEIN), the Mission Cancer within HORIZON EU Programme (PANCAID-101096309) and Fundación Ramón Areces. J.G.-A. was supported by Fundación Científica de la Asociación Española Contra el Cáncer (Predoctoral AECC 2021, PRDMA21602GARC). We thank to all members of H. Peinado’s laboratory and people/institutions involved for their dedication and contributions to the ideas discussed in this work.

## Extended Data Figure Legends

**Extended Data Fig. 1.**
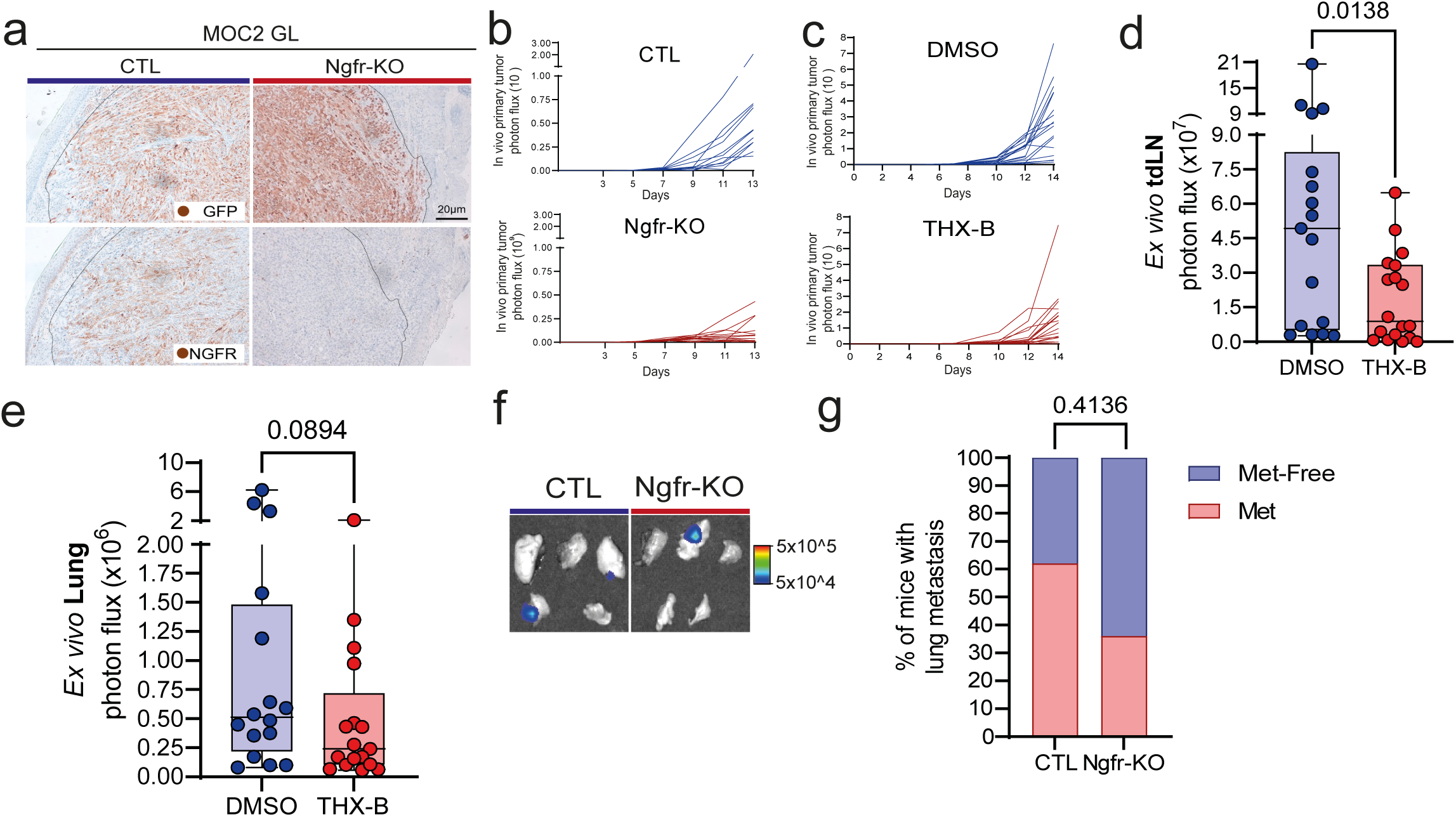
| Ngfr-KO validation, tumor growth impairment, and blockade of NGFR reduce spontaneous metastasis. a, Validation of MOC2 Ngfr-KO in vivo by IHC on orthotopic tumors. GFP was stained to differentiate tumor regions. Sample size: n = 5 mice per group. b, Individual mouse growth curves of MOC2 GL CTL and Ngfr-KO tumors, based on in vivo BLI at D3, 5, 7, 9, 11, and 13. Sample size: n = 11 for MOC2 GL CTL–bearing mice and n = 13 for MOC2 GL Ngfr-KO–bearing mice, pooled from two experiments. c, Individual mouse growth curves of MOC2 GL tumors treated with DMSO or THX-B. Sample size: n = 19 for DMSO-treated, and n = 18 for THX-B-treated mice, pooled from two independent experiments. d, e, Analysis of spontaneous metastasis after 0.05% DMSO or THX-B treatment of MOC2 GL tumor-bearing mice. Quantification of tdLN metastasis is shown in d, and lung metastasis is shown in e. Each dot represents an individual mouse; tdLN data reflects the combined signals from both cervical lymph nodes, and lung data reflects the combined signals from all five lung lobes. Statistical comparisons were performed using an unpaired t-test. Sample sizes: n = 17 DMSO and n = 18 THX-B for tdLN metastases, and n = 16 DMSO and n = 17 THX-B for lung metastases, pooled from two independent experiments. f, Representative BLI images of lung metastases from one mouse per group in an experimental metastasis model, in which mice were injected intravenously via the tail vein with 10⁵ MOC2 GL cells (CTL or Ngfr-KO) and metastatic burden in the lungs was assessed by ex vivo BLI at day 27. g, Quantification of lung metastases. P value was calculated using Fisher’s exact test. The percentages of mice in each category are shown, together with the corresponding P value. Sample size: n = 13 for CTL and n = 11 for Ngfr-KO, from two independent experiments.

**Extended Data Fig. 2.**
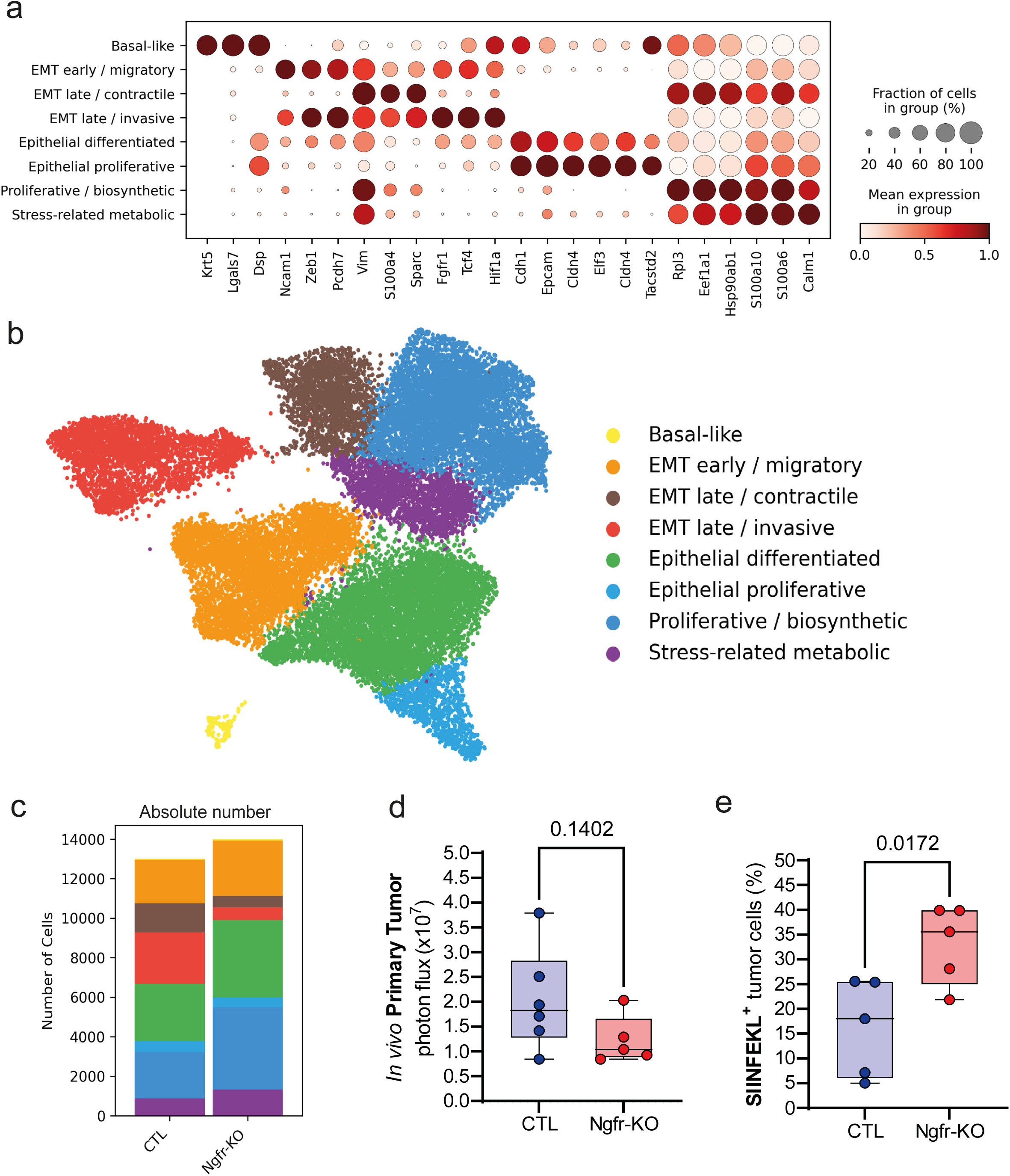
| MOC2 GL tumor cell clusters and evaluation of SIINFEKL expression in CTL and Ngfr-KO with similar tumor size using MOC2 GL. a, Dot plot showing the main genes used for cluster annotations, obtained through unbiased clustering using Scanpy. b, UMAP representation of tumor clusters across scRNA-seq samples from GFP^+^ tumor cells derived from MOC2 GL CTL or Ngfr-KO tumors. c, Stacked bar plot showing the absolute number of cells of each tumor cluster in MOC2 CTL or Ngfr-KO samples. d, Tumor growth quantification by BLI in MOC2 GL CTL and Ngfr-KO tumors. Each point represents an individual mouse. Statistical comparisons were performed using an unpaired t-test. e, Percentage of MOC2 GL tumor cells expressing SIINFEKL in the plasma membrane at D9. Statistical comparison was performed by an unpaired t-test. Sample size: n = 7 per group, single experiment.

**Extended Data Fig. 3.**
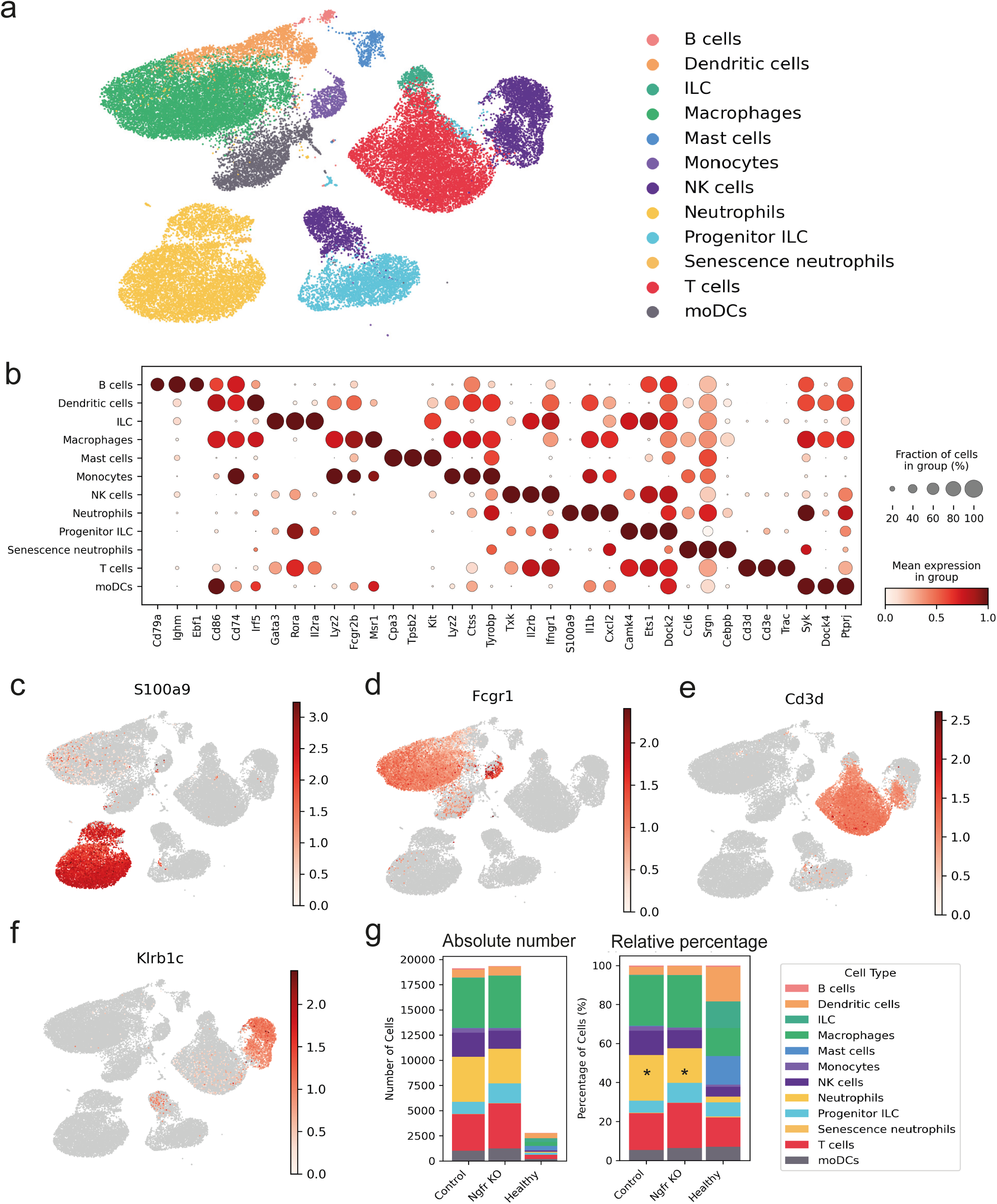
| Characterization of MOC2 GL immune cell clusters. a. UMAP representation of annotated immune populations derived from the scRNA-seq dataset of CD45+ cells from MOC2 GL tumors. b, Dot plot of classical markers corresponding to each immune population identified. c-f, UMAP of specific markers representative of the principal immune populations identified by scRNA-seq: S100a9 as a neutrophil marker is shown in c; Fcgr1 as a macrophage marker is shown in d; Cd3d as a T cell marker is shown in e; and Klrb1c as an NK cell marker is shown in f. g, Stacked bar plots showing the absolute number of cells (left) and the relative percentage (right) of each immune cell cluster in MOC2 GL CTL and Ngfr-KO tumors, together with healthy (non-tumor-bearing) tissue as a reference. Statistical significance (denoted by an asterisk) was determined using a two-step approach: (I) a contingency table test (Fisher’s exact test for cell counts <5 or chi-square test otherwise) with FDR adjustment (FDR < 0.05), and (II) biological relevance defined as an absolute percentage change greater than 5%.

**Extended Data Fig. 4.**
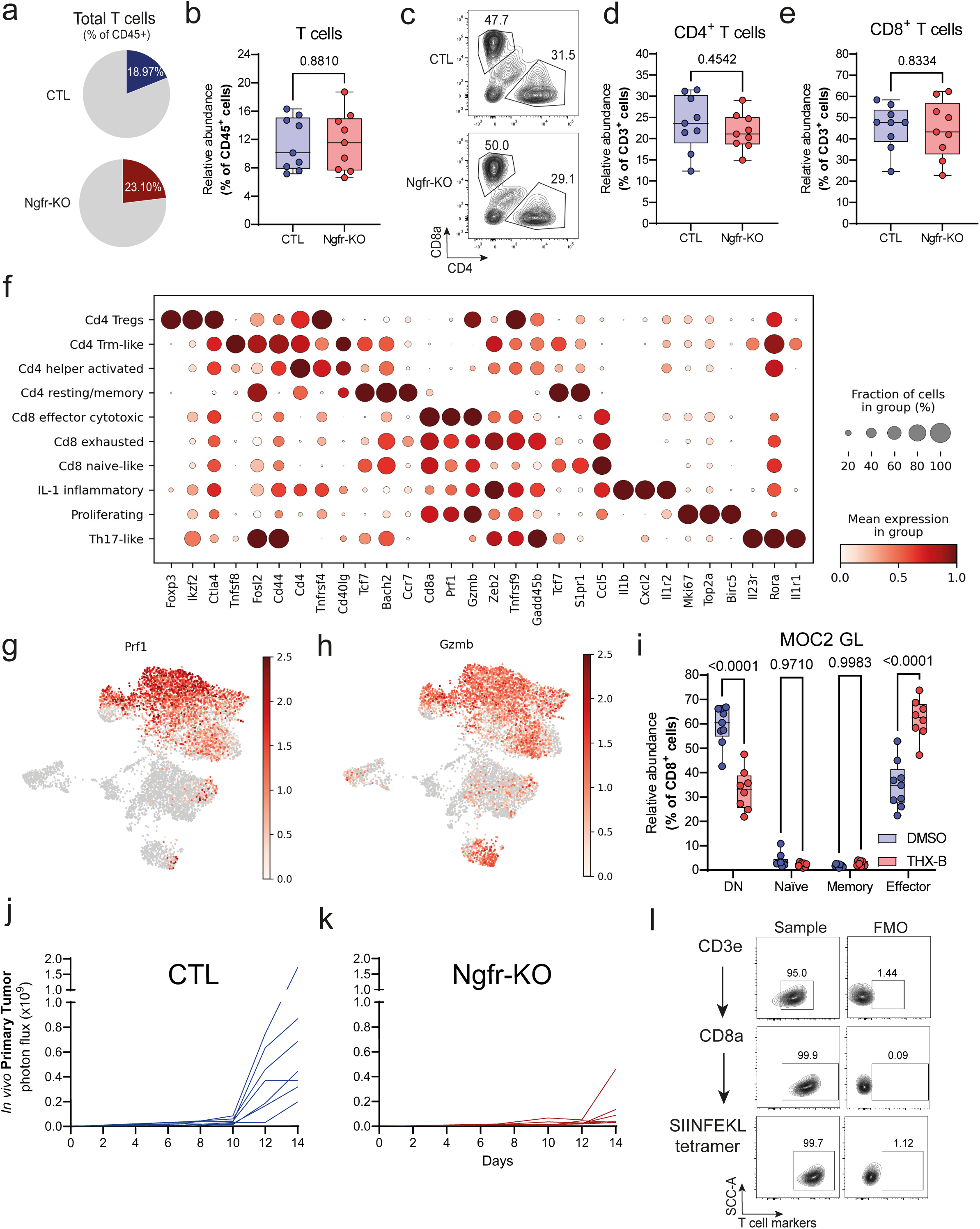
| NGFR does not affect general T cell populations but specifically modulates effector CD8+ T cells. a, Schematic representation of the percentage of T cells in the CD45^+^ scRNA-seq analysis from MOC2 GL CTL or Ngfr-KO tumors. Percentage relative to total CD45^+^ cells is shown. b, Relative abundance of T cells (CD45^+^CD3^+^) among total CD45^+^ cells, determined by flow cytometry. P values were calculated using an unpaired t-test. Sample size: n = 9, from three independent experiments. c, Representative density plot of MOC2 GL CTL or Ngfr-KO tumors showing the relative percentages of CD8a+ and CD4+ cells within the CD3^+^ population. d, e, Quantification of the relative abundance of CD4^+^ T cells (CD45^+^CD3^+^CD4^+^) and CD8^+^ T cells (CD45^+^CD3^+^CD8^+^) relative to total T cells (CD45^+^CD3^+^). The percentage of CD4+ cells is presented in d, and that of CD8+ cells in e. P values were calculated by an unpaired t-test. Sample size: n = 9, combining three independent experiments. f, Dot plot showing some of the most representative and differentially expressed genes in each T-cell subcluster identified in the scRNA-seq (three genes within the top 100 DEGs per cluster are represented). g, h, UMAP representations of Prf1 expression in g, and Gzmb expression in h, in T-cell subclusters (subpopulation assignment is shown in Fig. 3). i, Relative abundance of CD8+ T cell subtypes (DN, naïve, memory and effector) by flow cytometry in MOC2 GL tumors treated with 0.05% DMSO or THX-B (100 µg/mouse), assessed at D14. Statistical comparisons were performed using two-way ANOVA. Sample size: n = 9 for the DMSO group and n = 8 for the THX-B group, combined from two independent experiments. j, k, Longitudinal growth of MOC2 GL OVA-mCherry CTL or Ngfr-KO primary tumors based on *in vivo* BLI at D7, 10, 12, and 14. Individual mouse growth curves are shown. Sample size: n = 7 per group, pooled from two experiments. l, Density plots representing the expression of CD3e (top), CD8a (middle) and SIINFEKL-tetramer (bottom) in OT-I T cells derived from adoptive transfer experiments on MOC2 GL OVA-mCherry models. One representative sample (left) and the corresponding FMO (right) are represented.

**Extended Data Fig. 5.**
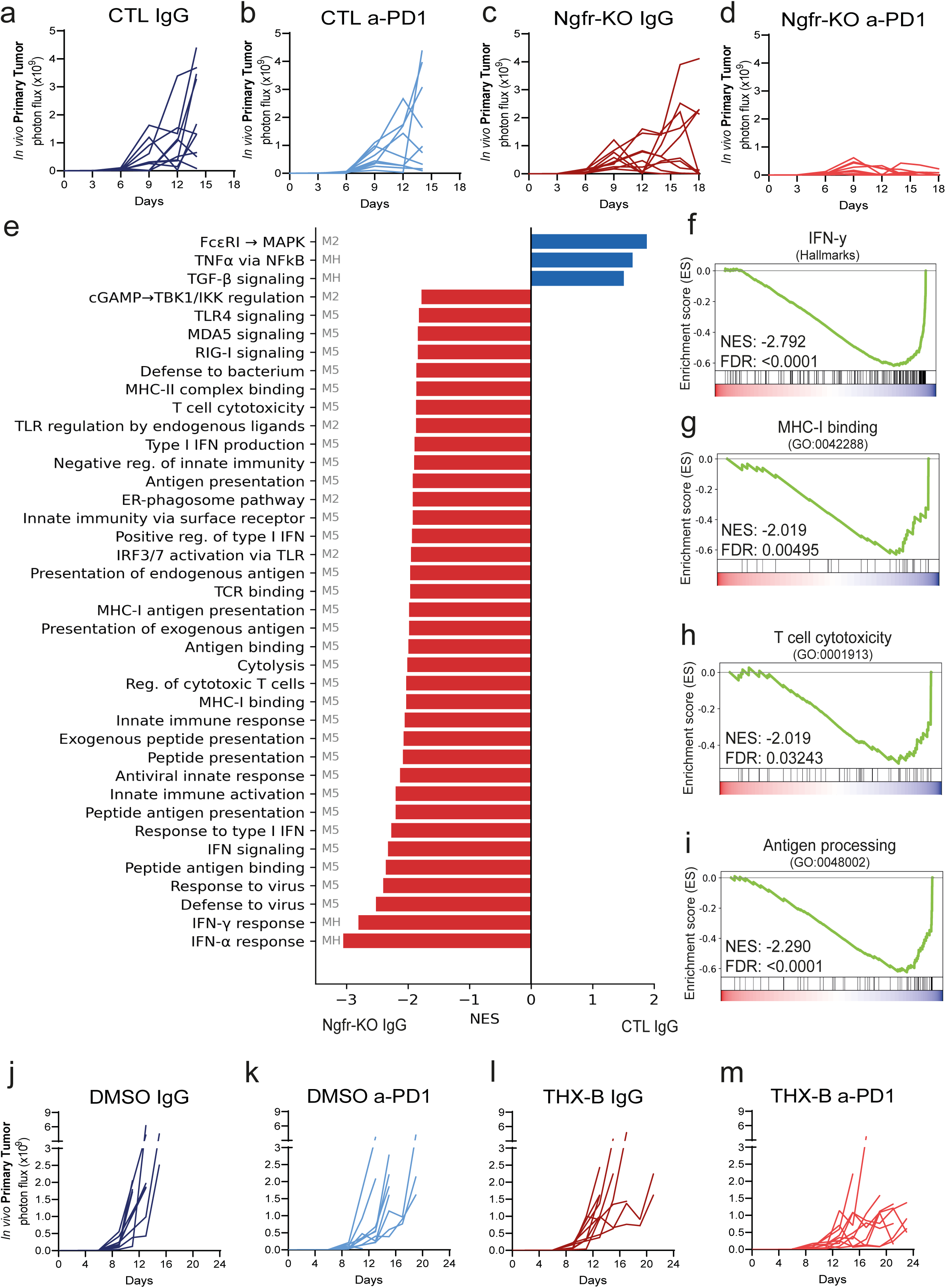
| Blocking NGFR enhances the response to anti-PD1 treatment, and its deletion promotes the activation of immune pathways. a–d, Individual tumor growth curves determined by BLI in mice bearing MOC2 GL CTL or Ngfr-KO tumors treated with IgG or anti-PD1. a, MOC2 GL CTL + IgG; b, MOC2 GL CTL + anti-PD1; c, MOC2 GL Ngfr-KO + IgG; d, MOC2 GL Ngfr-KO + anti-PD1. Each line represents an individual mouse. Sample sizes: CTL IgG (n = 10), CTL anti-PD1 (n = 9), Ngfr-KO IgG (n = 10), Ngfr-KO anti-PD1 (n = 10), from a single experiment. e, GSEA of all significantly upregulated immune pathways (FDR < 0.05) when comparing IgG-treated MOC2 GL CTL vs. Ngfr-KO tumors. Mouse-specific datasets from Hallmarks (MH), Reactome (M2), and Gene Ontology (M5) were combined into a single graph, with the source indicated to the left of each pathway name. FDR values were calculated using the Benjamini-Hochberg method to correct for multiple comparisons. f-i, Representative examples of the immune-related pathways enriched in Ngfr-KO tumors are shown, including IFN-γ response (Hallmark) in panel f, MHC class I protein binding (GO:0042288) in panel g, T cell–mediated cytotoxicity (GO:0001913) in panel h, and antigen processing and presentation of peptide antigen (GO:0048002) in panel i. j–m, Individual tumor growth curves determined by BLI in mice with MOC2 GL tumors treated with: j, DMSO + IgG; k, DMSO + anti-PD1; l, THX-B + IgG; m, THX-B + anti-PD1. Each line represents an individual mouse. Sample sizes: n = 10 mice per group, pooled from two independent experiments.

**Extended Data Fig. 6.**
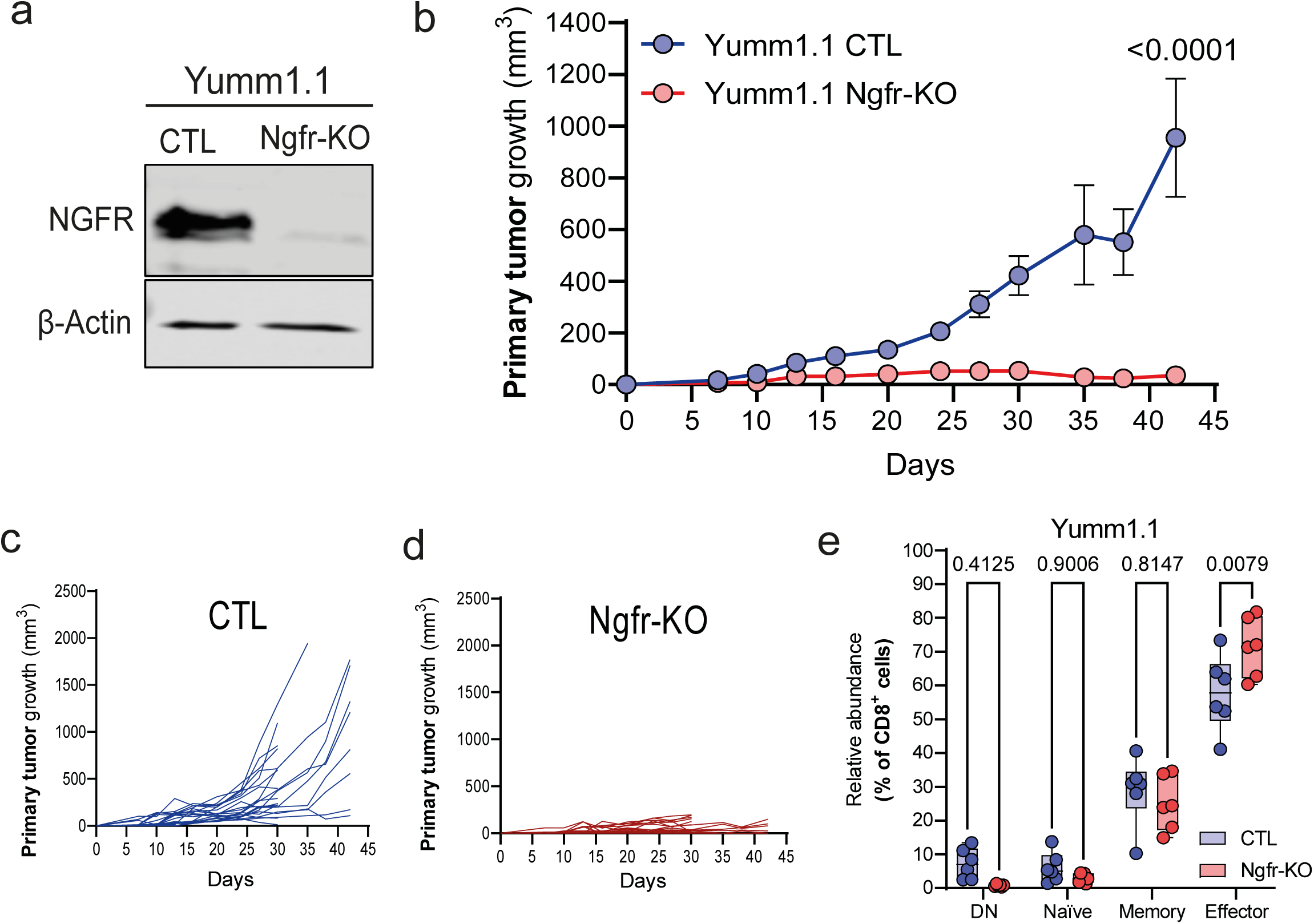
| Ngfr-KO validation, tumor growth impairment in Yumm1.1 tumors, and expansion of effector CD8⁺ T cells. a, Ngfr-KO validation in the Yumm1.1 cell line. A representative immunoblot is shown (n = 3). b–d, Tumor growth curves of Yumm1.1 CTL and Ngfr-KO groups, represented as the mean ± SEM in b, and as individual tumor volumes per mouse in c and d. Statistical significance across time points was determined using two-way ANOVA, with the P value corresponding to the endpoint shown in b. Sample size: n = 11 mice per group from two independent experiments (CTL: 22 tumors; Ngfr-KO: 21 tumors). Individual tumors were treated as the unit of analysis, consistent with common practice in bilateral flank tumor models; each mouse contributed one tumor per flank. e, Relative abundance of CD8+ T cell subtypes by flow cytometry in Yumm1.1 CTL or Ngfr-KO tumors (D40). P values were calculated by two-way ANOVA. Sample size: n = 6 tumors per group in two independent experiments.

**Extended Data Fig. 7.**
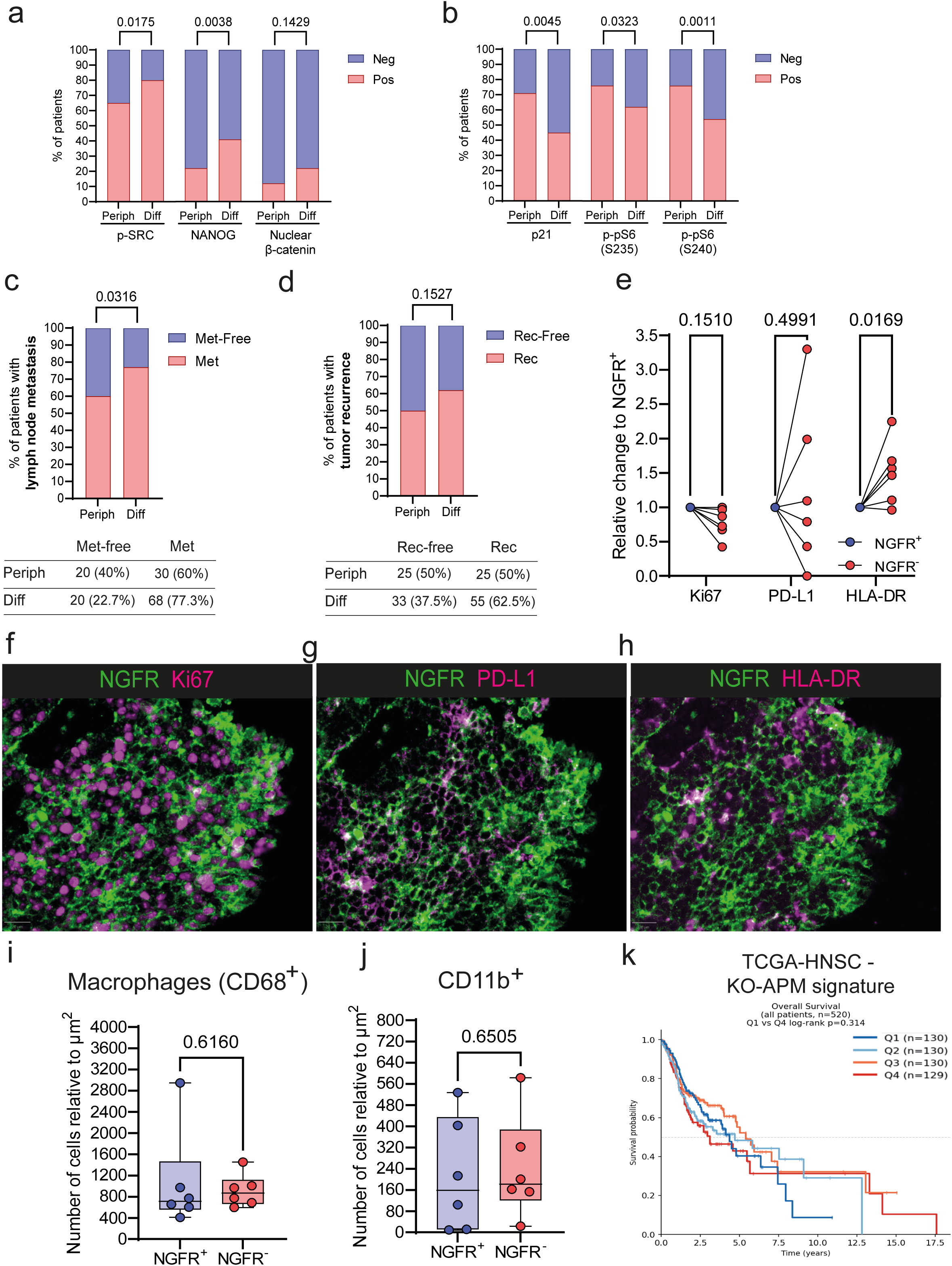
| Association of NGFR with aggressive clinical features, tumor markers, immune infiltrates, and antigen-presentation signature in human HNSCC tumors. a, b, Association between NGFR expression patterns and prognostic markers. Panel a shows markers associated with poor prognosis (p-SRC, NANOG, and nuclear β-catenin), whereas panel b shows markers associated with good prognosis (p21, p-pS6 S235, and p-pS6 S240). The percentage of patients positive and negative for each marker is plotted. P values were calculated using Fisher’s exact test, comparing the total number of patients in each group. c, d, Contingency plots illustrate the association between NGFR expression patterns and clinical features. Panel c shows the relationship between NGFR expression patterns and the presence of lymph node metastasis, whereas panel d shows the association with tumor recurrence. P values were calculated using Fisher’s exact test. The percentages of patients in each category are shown, together with the corresponding P values. A summary table with the number and percentage of patients in each group is also included. e, Quantification of relative marker expression in PanCK⁺NGFR⁺ versus PanCK⁺NGFR⁻ tumor cells for Ki67, PD-L1, and HLA-DR in HNSCC. Data are shown for n = 6 patients from two independent experiments. P values were calculated from per-tumor percentages using a paired two-sided t-test, pairing values from the same tumors. f– h, Representative high-plex immunofluorescence images from the same tumor region of one HNSCC patient showing NGFR (green) co-staining with different markers (magenta): Ki67 (f), PD-L1 (g), and HLA-DR (h). i, Relative frequency of macrophages (CD68⁺) in PanCK⁺NGFR⁺ and PanCK⁺NGFR⁻ regions, normalized to the total area in each case. j, Relative frequency of CD11b⁺ myeloid cells in PanCK⁺NGFR⁺ and PanCK⁺NGFR⁻ regions, normalized to the total area in each case. Data in i–j are from n = 6 patients from two independent experiments; P values were calculated using paired two-sided t-tests. k, Kaplan–Meier overall survival analysis of HNSCC patients from TCGA stratified by an NGFR-KO–derived antigen-presentation signature by quartiles of expression. The P value was calculated using the log-rank test.

